# Transcription factors SP5 and SP8 drive primary cilia formation

**DOI:** 10.1101/2025.06.03.657415

**Authors:** Yinwen Liang, Richard Koche, Ravindra B. Chalamalasetty, Daniel N. Stephen, Mark W. Kennedy, Zhimin Lao, Yunong Pang, Ying-Yi Kuo, Moonsup Lee, Francisco Pereira Lobo, Xiaofeng Huang, Anna-Katerina Hadjantonakis, Terry P. Yamaguchi, Kathryn V. Anderson, Alexandra L. Joyner

## Abstract

While specific transcription factors are known to regulate cell fate decisions, the degree to which they can stimulate formation of specific cell organelles is less clear. We used a multi-omics comparison of the transcriptomes of ciliated and non-ciliated embryonic cells to identify transcription factors upregulated in ciliated cells, and conditional genetics in mouse embryos and stem cells to demonstrate that SP5/8 regulate cilia formation and gene expression. In *Sp5/8* mutant embryos primary and motile cilia are shorter than normal and reduced in number across cell types, contributing to situs inversus and hydrocephalus. Moreover, expression of SP8 is sufficient to induce primary cilia in unciliated cells. This work opens new avenues for studying cilia assembly using stem cell models and offers new insights into human ciliopathies.

## Main Text

Cilia are microtubule-based organelles that project from most cell types, including proliferating progenitors, and serve critical functions in motility, sensation, and signal transduction (*1, 2*). Accordingly, genetic mutations that disrupt cilia cause developmental abnormalities termed ciliopathies that include hydrocephalus, situs inversus, polycystic kidney disease, and craniofacial defects (*1, 3, 4*). There are two types of cilia: primary and motile cilia based on their microtubule cytoskeleton (axoneme) structure. Most primary cilia are non-motile, solitary, and critical for signal transduction, particularly Hedgehog signaling, and can have sensory functions (*1, 2, 4*). Motile cilia provide cell motility or fluid flow and are present in multiple copies on terminally differentiated cells (*4*). There are hundreds of cilia proteins, and most are shared by primary and motile cilia (*5–7*). Two TF families (Forkhead and Regulatory factor X) have been found to enhance formation of sensory/primary cilia in a cell type specific manner and drive motile cilia formation across cell types and species (*8–18*). Despite the importance of primary cilia, the transcriptional control of genes required for primary cilia formation has not been delineated.

During mouse embryonic development, primary cilia form progressively starting at embryonic day 6 (E6.0) in pluripotent epiblast cells (*19*). Mouse embryonic stem cells (ESCs) and to a greater extent their derivative epiblast stem cells (EpiSCs) have cilia, as do human pluripotent stem cells (*19, 20*). Interestingly, while cells derived from the epiblast, including extraembryonic mesoderm, form cilia, the cells of the extraembryonic trophectoderm and the yolk sac visceral endoderm (YsVE) do not form cilia (*19*). YsVE have a high level of cilia disassembly protein activity, however, this does not explain the lack of primary cilia since drug inhibition of the cilia disassembly factors only results in rare YsVE cells forming primary cilia (*19*). This result raised the question of whether some cilia genes are poorly expressed in YsVE because the transcriptional activators for the genes are only present in ciliated cell types.

### scRNA-seq analysis reveals broad upregulation of cilia genes in ciliated cells versus YsVE

During development the visceral endoderm (VE) generates cells that populate the yolk sac (YsVE) and the embryo (embryonic VE or emVE) (Fig. 1A). The latter initially overlie the epiblast and then intercalate with embryo-derived definitive endoderm (DE) and the two cell types form the gut endoderm during E7.0-7.5 (*21*). At around E5.0, emVE becomes morphologically and molecularly distinct from YsVE (*22*). We asked whether both VE- and DE-derivatives in the gut endoderm have cilia by employing two VE-specific reporter lines, *Afp-GFP* (*23*) and a combination of *Ttr-Cre* (*24*) (VE-specific driver) and a tdTomato (tdT) Cre reporter *R26^lsl-tdT^* (*25*) (Fig. 1B, fig. S1A, B). We confirmed a lack of cilia in YsVE using transmission electron microscopy and found that while the centrioles were apically localized, they failed to dock to the apical membrane and recruit ciliary vesicles necessary for cilia formation (fig. S1C, D). Strikingly, the emVE began to form cilia at around E7.0, and by E8.75 when the gut endoderm had formed, most gut VE cells (GutVE) had cilia (Fig. 1C-E, G, fig. S1A, B). Cilia formation in epiblast-derived DE and mesoderm cells peaked earlier at E6.5 (Fig. 1G). Moreover, cilia formation in the emVE lineage was concurrent with their intercalation with DE cells during E7.0 and E7.5 (fig. S1E). Furthermore, intercalation with DE cells seems to be required for cilia formation in emVE since in mutants in which mesoderm and DE do not form (*Fgf8^-/-^*) or DE forms but fails to fully interact with emVE (*Epi-Strip1*) emVE cells do not form cilia (fig. S1F-K) (*26, 27*). Thus, intercalation of emVE with DE could contribute to a DE-like transcriptional signature that includes cilia genes.

**Fig. 1.**
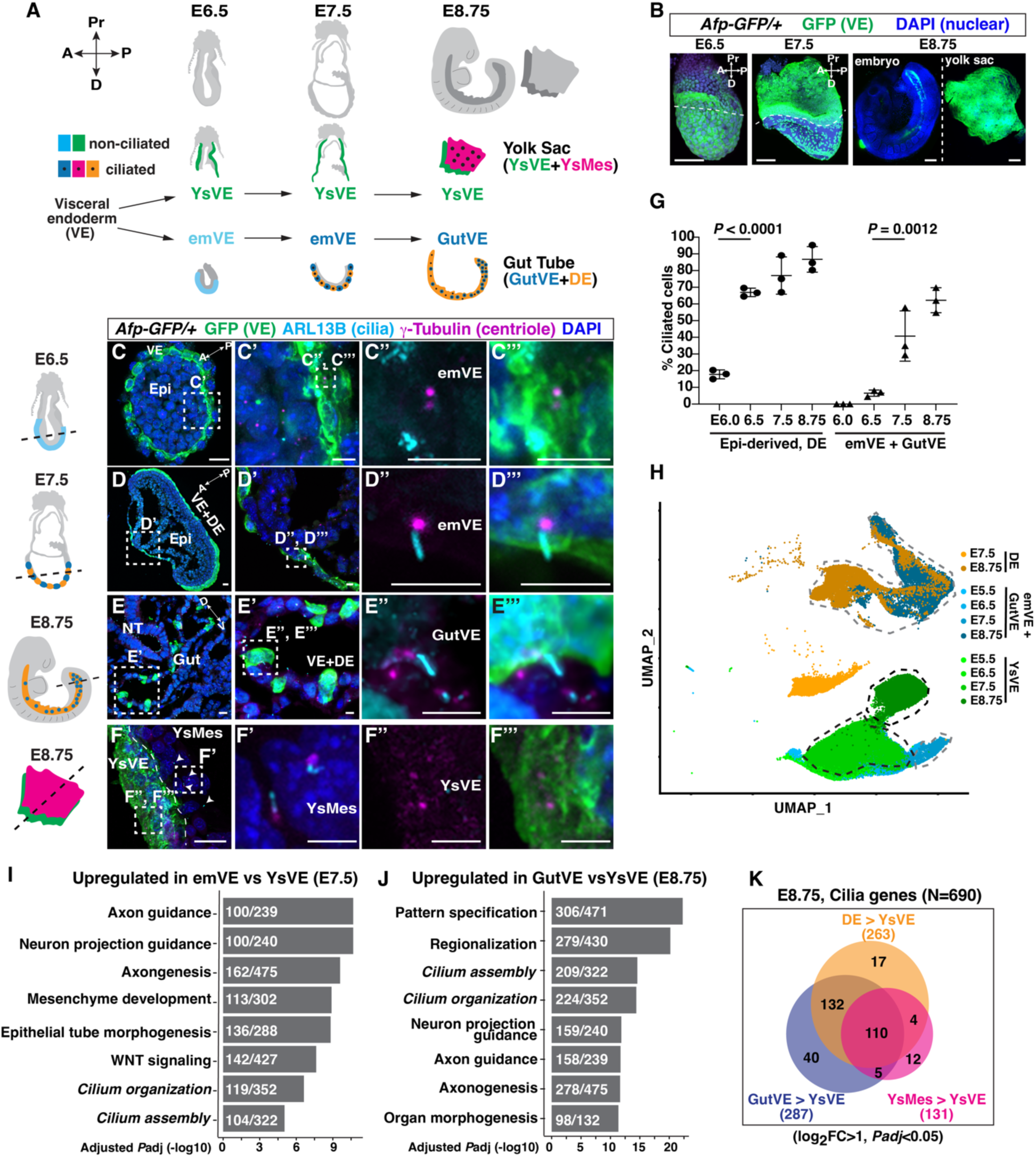
VE-derived emVE cells form cilia during intercalation with DE to form gut. (**A**) Schematic showing that the extraembryonic visceral endoderm (VE) lineage generates yolk sac VE (YsVE, green) and embryonic VE (emVE, cyan). emVE intercalates with embryo-derived definitive endoderm (DE, orange) of the gut to become gut VE (GutVE, blue). Ys also contains embryo-derived mesoderm (YsMes, pink). Black dots, ciliated cells. The schematics of embryos in (A) and beside (C-E) were modified from (*21*). (**B**) Wholemount images of *Afp-GFP/+* embryos. Extra-embryonic tissue, above dashed line (E6.5, E7.5); Ys, right (E8.75); Pr, proximal; D, distal. (**C**-**F)** Immunofluorescent (IF) staining of transverse sections of *Afp-GFP/+* embryos. Schematics indicate section level. Arrowheads, ciliated YsMes (F); D, dorsal; V, ventral; epi, epiblast; A, anterior; P, posterior. Scale bar, 1mm (B); 20μm low magnification, 5μm insets (C-F). (**G)** Quantification of the percentage ciliated emVE and GutVE (E6.0-E8.75), epiblast-derived cells (E6.0-E7.5), and DE (E8.75). Statistical analysis: one-way ANOVA (N=3 embryos/stage). (**H**) Uniform Manifold Approximation and Projection (UMAP) plot showing clustering of cells (scRNA-seq from (*15*)). (**I**, **J**) Bar plots of the top 6 upregulated biological processes and cilium-related terms in E7.5 emVE (I), the top 8 upregulated biological processes in E8.75 GutVE (J) compared to YsVE. Input gene cutoff log_2_Fold Change>1; *Padj*<0.05. Gene sets are ranked by *Padj*. The numbers in the boxes indicate the number of upregulated genes per total gene number per term. See also table S2, S3. (**K**) Venn diagram showing the overlap of upregulated cilia genes in GutVE, DE, or YsMes compared to YsVE at E8.75 (N=690 cilia genes, table S1).

To determine if some cilia genes are expressed at a higher level in ciliated cells, we profiled the expression of cilia genes in three ciliated cell types compared to unciliated YsVE. We chose a recently curated list of cilia genes that includes ones that regulate primary and motile cilia formation, and contribute to structure, function, and the transport machinery (N=690, table S1) (*7*). Some cilia proteins have additional functions outside the cilium and therefore are expressed in most cell types (*28, 29*). We first asked whether within the VE-lineage a subset of cilia genes is expressed at a higher level in GutVE (ciliated) cells compared with YsVE (unciliated). We subsetted the E5.5-E8.75 DE, emVE, GutVE, and YsVE cells from a single cell RNA-sequencing (scRNA-seq) data set of the early endoderm (*21*), and as expected the four cell types were represented as clusters by age (Fig. 1H). Differential gene expression analysis between GutVE and YsVE at each age followed by Gene Ontology (GO) enrichment analysis identified cilium organization and cilia assembly terms as significantly upregulated at E7.5 and were in the top 4 signatures at E8.75. (*Padj* < 0.05; Fig. 1I, J, table S2, 3). Among the cilia genes, 287 genes were significantly upregulated at least 2-fold at E8.75, 157 at E7.5, and 65 at E6.5 in GutVE compared to YsVE (log_2_FC [Fold Change]>1, *Padj*<0.05, fig. S2A, B, table S4). 304 genes were significantly upregulated more than 2-fold in at least one stage, with 50 genes upregulated at all three stages (fig. S2B, table S4).

To investigate whether these cilia genes are upregulated in other cell types with primary cilia, we compared the transcriptome of YsVE with its neighbor cells, yolk sac mesoderm (YsMes), or with gut DE. We performed scRNA-seq on E8.75 yolk sac tissue, and cluster analysis identified the expected five cell types (fig. S2C, table S5). Differential gene expression analysis revealed that cilium organization and cilia assembly GO terms were upregulated in E8.75 YsMes compared to YsVE (fig. S2D, E, table S6, 7). These Go terms were also upregulated after comparing E8.75 DE with YsVE (fig. S2F, G, table S8, 9). Curiously, both primary cilia and some motile cilia genes were enhanced in all three ciliated cell types. Interestingly, differential gene expression analysis showed that most of the 287 cilia genes upregulated in E8.75 GutVE compared to YsVE were also upregulated in DE or YsMes compared to YsVE (242/263 genes upregulated in DE, 115/131 upregulated in YsMes), with 110 genes upregulated in all three ciliated cell types at E8.75 (Fig. 1K, fig. S3A, table S10). Thus, we identified a set of 110 cilia genes that have shared upregulation in three ciliated cell types compared to unciliated YsVE at E8.75. Expression across lineages of these shared cilia genes appears to be conserved in human embryos, as the average expression was significantly downregulated in yolk sac endoderm compared to ciliated embryonic cells in human and mouse, whereas the remaining cilia genes were not downregulated (fig. S3B) (*30*). This finding supports our idea that a transcriptional activator promotes cilia gene expression.

### Chromatin accessibility analysis identifies SP/KLF family TFs as binding cilia genes

We next asked if there are TFs that bind to open chromatin in the shared cilia genes (Fig. 2A) by performing **a**ssay for **t**ransposase-**a**ccessible **c**hromatin with sequencing (ATAC-seq) on GutVE and YsMes compared to unciliated YsVE at E8.75 embryos (Fig. 2B). Upon peak calling, cilia genes were enriched in the enhanced open chromatin in GutVE and YsMes compared to YsVE (fig. S4, table S11, 12). 267 of the 690 cilia genes had enhancement of open chromatin in ciliated cells compared to YsVE (log_2_FC>0, *Padj*<0.05, table S13). Strikingly, the ATAC-seq analysis showed that of the 110 shared cilia genes, 70 had enhancement of open chromatin in promoter or enhancer regions (N=203 peaks, table S14). Moreover, motif enrichment analysis on these 203 peaks showed that DNA motifs specific to Krüppel-like factor and Specificity protein (KLF/SP) family TFs were significantly enriched (Fig. 2C, table S15). DNA binding motifs for CTCF, CTCFL, Fork head family (FOXP1, FOXA2/3, FOXF1, FOXO3) and TEAD1 were enriched to a lesser extent. Motif analysis of all cilia genes (N=690) with enhanced chromatin regions (N=586 peaks of 267 genes, table S16) also identified the same TF families, except TEAD1, and the motile cilia TFs RFX2, RFX3 were included but in a low percentage of peaks (∼6%) (fig. S5A).

**Fig. 2.**
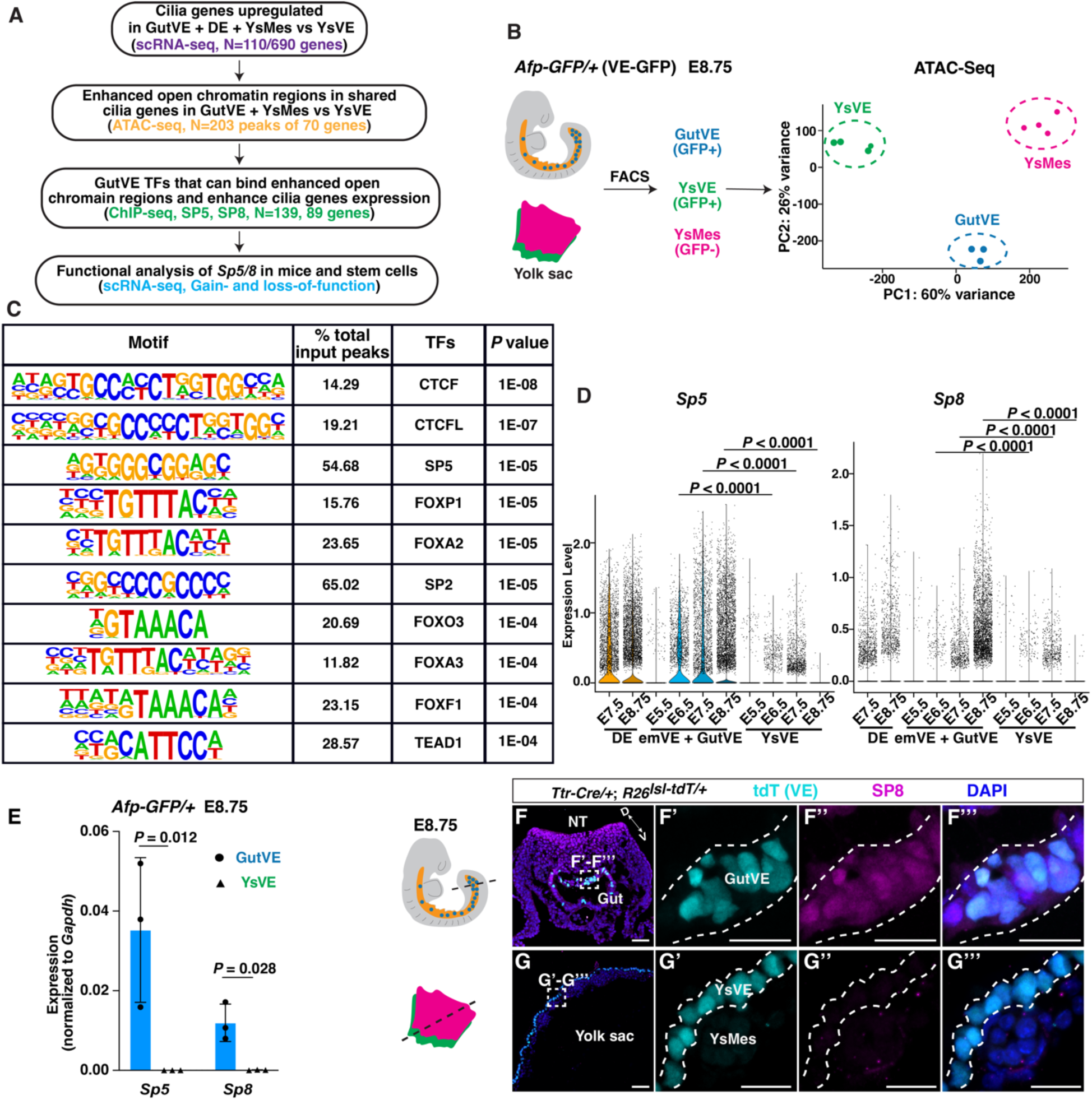
Chromatin accessibility analysis of core ciliome genes in GutVE and YsM compared to YsVE identifies SP/KLF TF families. (**A**) Study design, scRNA-seq and ATAC-seq in E8.75 embryo and ChIP-seq in ES cell lines. (**B)** Schematic diagram showing ATAC-seq sample preparation (left) and scatter plot (right) of sample clustering from principal component analysis (PCA) of GutVE, YsMes, and YsVE (N=3, 4, 4 samples). See also tables S11-15. (**C)** Enrichment of transcription factor recognition sequences in 203 differential ATAC-seq peaks in 70 of the shared cilia genes in GutVE and YsMes versus YsVE. The list is ranked based on *P* value. See also table S15. (**D)** Violin plot showing the log_10_ CPM (counts per million) expression levels of *Sp5* and *Sp8* from scRNA-seq data. See also table S17. (**E**) qRT-PCR quantification of *Sp5*, *Sp8* mRNA expression normalized to *Gapdh*. (**F**, **G)** IF staining of transverse sections for SP8, tdT (VE lineage cells), and DAPI in the gut tube (F) and yolk sac (G) of E8.75 *Ttr-Cre/+*; *R26^lsl-tdT/+^* embryos. (F’-F’’’, G’-G’’’) Higher magnification images of insets in (F, G). D, dorsal; V, ventral. Scale bar, 50μm, insets 20μm. The schematic of the embryo in (B) and beside (F) was modified from (*21*). Statistical analyses: Wilcoxon rank-sum test (D), unpaired *t*-test (E).

KLF/SP family TFs bind to GC-box or GT-box elements in promoters and enhancers and play many roles during development (*31*). Importantly, among the 26 SP/KLF family genes (*Sp1-9*, *Klf1-17*), only two (*Sp5*, *Sp8*) were differentially expressed in emVE+GutVE compared to YsVE at E6.5, E7.5 and E8.75 (Fig. 2D, fig. S5B, C, table S17). Moreover, *Sp5* and *Sp8* expression increased over the time when cilia form (Fig. 2D). *Klf3* and *Klf12* expression were significantly increased only at E8.75. Furthermore, *Klf12* expression was very low in all cell types and KLF3 protein was previously shown to only be detected before E3.5 (fig. S5D) (*32*). Among Fork head family genes, *Foxp1* was the only gene upregulated in GutVE compared to YsVE at the three ages and *Foxj1* was low but slightly upregulated in GutVE at E8.75 (fig. S5E). We focused our functional analysis on *Sp5* and *Sp8* based on their gene expression, high percentage of motif enrichment and being novel candidates to regulate cilia.

SP5/8 mediate WNT/b-catenin signaling in multiple cell types and are expressed in the E8.5 gut endoderm where they promote hindgut extension and colon formation (*33–36*). To determine if the genes are expressed in the GutVE compartment of gut endoderm, we performed qRT-PCR analysis on E8.75 GutVE and YsVE and found *Sp5/8* were expressed in GutVE and not YsVE (Fig. 2E). Immunofluorescence analysis of tissue sections confirmed that SP8 protein is present in gut endoderm, including GutVE, and is low in YsMes and absent from the yolk sac at E7.5 and E8.75 (Fig. 2F, G, fig. S5F, G).

In summary, a screen for motif enrichment in open chromatin regions near cilia genes identified SP5 and SP8 as strong candidates to drive expression of a group of genes associated with cilia formation and function.

### SP5/8 bind to and are required for expression of a subset of cilia genes in ES cell-derived gastruloids

Based on previous SP5-**Ch**romatin **i**mmuno**p**recipitation followed by **seq**uencing analysis (ChIP-seq) in mouse embryoid bodies (GSE72989) (*35*), 139/690 cilia genes were bound by SP5 (N=202 binding sites) (table S17). ChIP-seq was performed to identify genes bound by SP8 in an ES cell line with doxycycline (Dox) inducible FLAG-tagged SP8 (*i-Sp8-3xFlag*) (*35*) (Fig. 3A, fig. S6A-C, table S18). EpiSCs were generated for ChIP-seq because they are highly ciliated compared to ESCs (82.2% vs 18.9%). FLAG ChIP-seq data analysis of Dox-induced *i-Sp8-3xFlag* cells (SP8-FLAG) versus control (no Dox) revealed that SP8 binds to 89/690 cilia genes (N=147 binding sites), 44 of which are shared with SP5 (Fig. 3B). In total, 187 cilia genes are bound by SP5 and/or SP8. SP8 preferentially binds to promoter regions (63.19%, N=78/89), while SP5 binds to promoter regions to a lesser extent (21.78%, N=40/139) (Fig. 3B, 3C, table S19, 20). The composite gene expression for SP5/8 bound cilia genes was significantly higher in GutVE compared to YsVE, and 100/187 genes were expressed at least 2-fold higher in GutVE, DE or YsMes compared to YsVE (Fig. 3D, table S21). The enhanced binding of SP5/8 preferentially occurred in regions of cilia gene open chromatin (Fig. 3E, fig. S6D, S6E). Interestingly, the motile cilia-related TF genes, *Foxj1* and *Rfx2/3/7* were bound by SP5/8 (fig. S7A). Like *Foxj1*, *Rfx2/3* were expressed at a higher level in ciliated cells than YsVE only at E8.75 (fig. S5C). We also found that among the 564 potential SP5/8 binding sites in cilia genes 98 also have RFX2/3 binding sites (N=214) within 500 bp (fig. S7B).

**Fig. 3.**
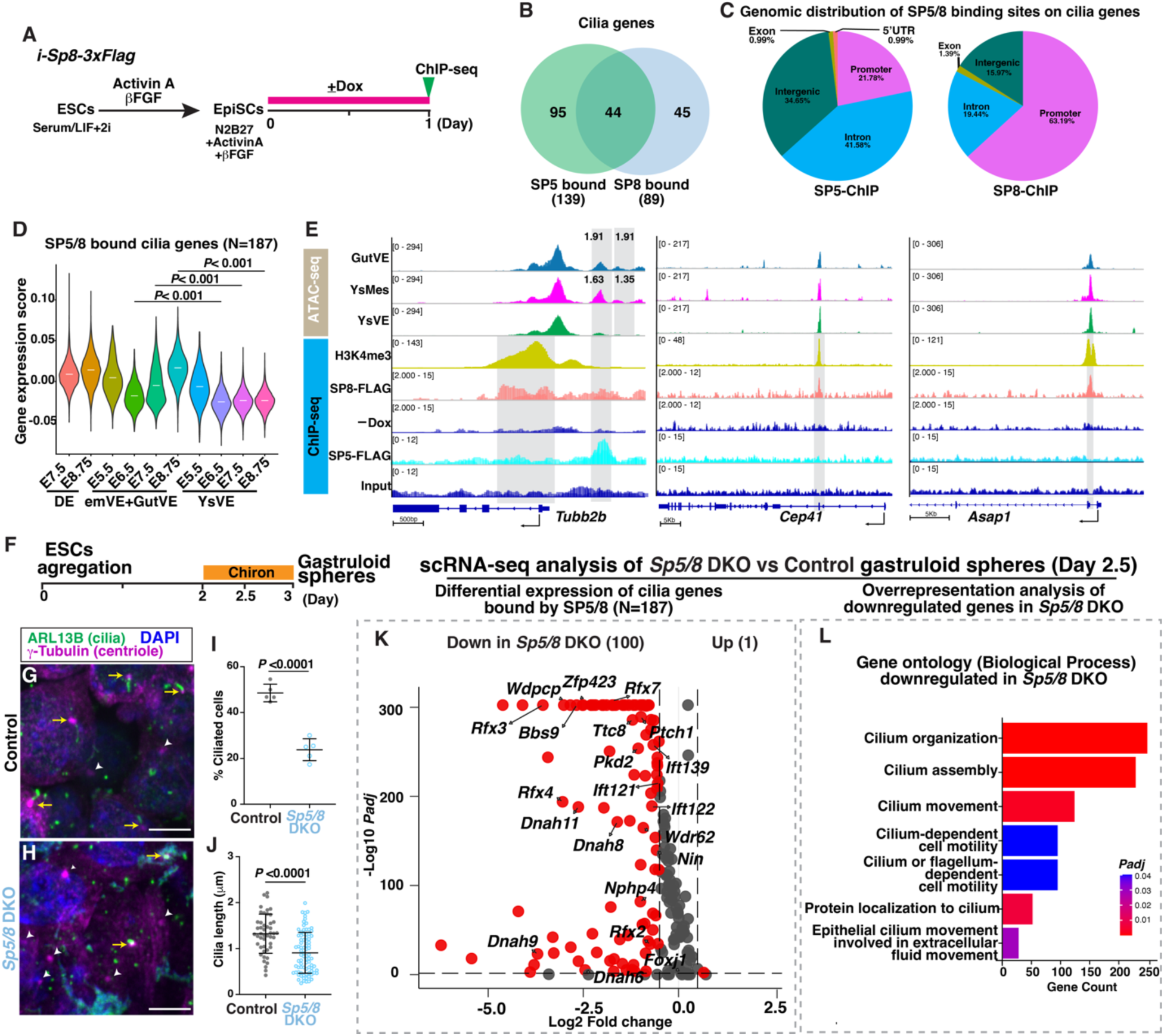
SP5/8 bind and activate cilia genes. (**A**) Experimental design. (**B**) Venn diagram showing the overlap of cilia genes bound by SP5 and SP8. See also tables S19, S20. (**C**) Gene distribution of SP5/8 binding sites in cilia genes. **(D)** Violin plots showing composite expression levels of SP5/8 bound cilia genes (N=187) in DE, GutVE, and YsVE at E5.5-E8.75. **(E)** ATAC-seq plots of cilia genes in GutVE, YsMes, YsVE and ChIP-seq tracks of H3K4me3 and SP8-FLAG in EpiSCs and SP5-FLAG in embryoid bodies. Grey boxes indicate significance of the peak comparisons of ATAC-seq and ChIP-seq data. The number above ATAC-seq tracks represents fold change in a single 500 bp peak with *Padj*<0.05. Y-axis scale was chosen to optimize the visualization of peaks for each sample. (**F**) Schematic showing derivation of gastruloid spheres from ESCs. (**G**, **H**) IF staining of WT and *Sp5/8* DKO ESCs derived day 2.5 gastruloid spheres. Yellow arrows indicate ciliated cells; white arrow heads indicate unciliated cells. (**I**, **J**) Quantification of the percentage of ciliated cells (I) and cilia length (J) in WT and *Sp5/8* DKO gastruloid spheres (N=5 technical replicates/genotype). (**K**) Volcano plot showing the differential expression of SP5/8 bound cilia genes (N=187) in *Sp5/8* DKO vs WT gastruloid spheres (table S22). (**L**) Overrepresentation analysis of downregulated genes in *Sp5/8* DKO vs WT gastruloid spheres showing the enrichment of cilia-related gene sets (table S23). Statistical analyses: Wilcoxon rank-sum test (D), unpaired *t*-test (I, J). Scale bars, 5μm (G, H).

We next tested whether *Sp5/8* are required for expression of cilia genes using gastruloid spheres derived from ESCs (*37, 38*). In the gastruloids derived from normal controls and *Sp5/8* double knockout (DKO) ESCs, the percentage of cells with cilia was decreased by half (23.8% in *Sp5/8* DKO vs 48.5% in control) and cilia length was also significantly reduced (0.9µm vs 1.3µm) (Fig 3.G-J). We then applied scRNA-seq analysis to control and *Sp5/8* DKO gastruloids and performed differential gene expression analysis. Strikingly, the majority of SP5/8 bound cilia genes were downregulated in the *Sp5/8* DKO gastruloids (100/187, table S22), including genes that encode IFT, BBS, NPHP, RFX2/3/7 and motile cilia structure proteins (Fig. 3K). Cilium-related gene sets were highly enriched in the downregulated genes in *Sp5/8* DKO gastruloids (Fig. 3L, table S23). Thus, SP5/8 play an important role in maintaining cilia gene expression and cilia formation in cells in culture.

### *Sp5/8* are required for cilia formation across embryonic cell types

Next, we tested whether *Sp5/8* are required for cilia formation in the developing mouse embryo. *Sp8* null mutants are lethal around E14, whereas *Sp5* mutants are viable with a kinked tail (*33, 34, 39*). We examined the VE lineage by generating mice with an *Sp5* null allele and VE-lineage conditional knock-out (cKO) of *Sp8* using *Ttr-Cre* and including a reporter allele to track the VE-lineage (*VE*-*Sp5/8* cKO) (*33*). SP8 protein was not detected in the GutVE of *VE*-*Sp5/8* cKO E8.75 embryos (fig. S8A, B). Importantly, the GutVE of *VE*-*Sp5/8* cKO embryos had a significant reduction in the frequency of ciliated cells compared to controls (39.7% of tdT+ gut cells in mutants; 62.8% in littermate controls), as well as a slight decrease in cilia length (1.37µm in mutants vs 1.55µm controls, *P*=0.0237, Fig. 4A-D). Therefore, SP5/8 play a role in generating cilia and promoting their normal length in the GutVE.

**Fig. 4.**
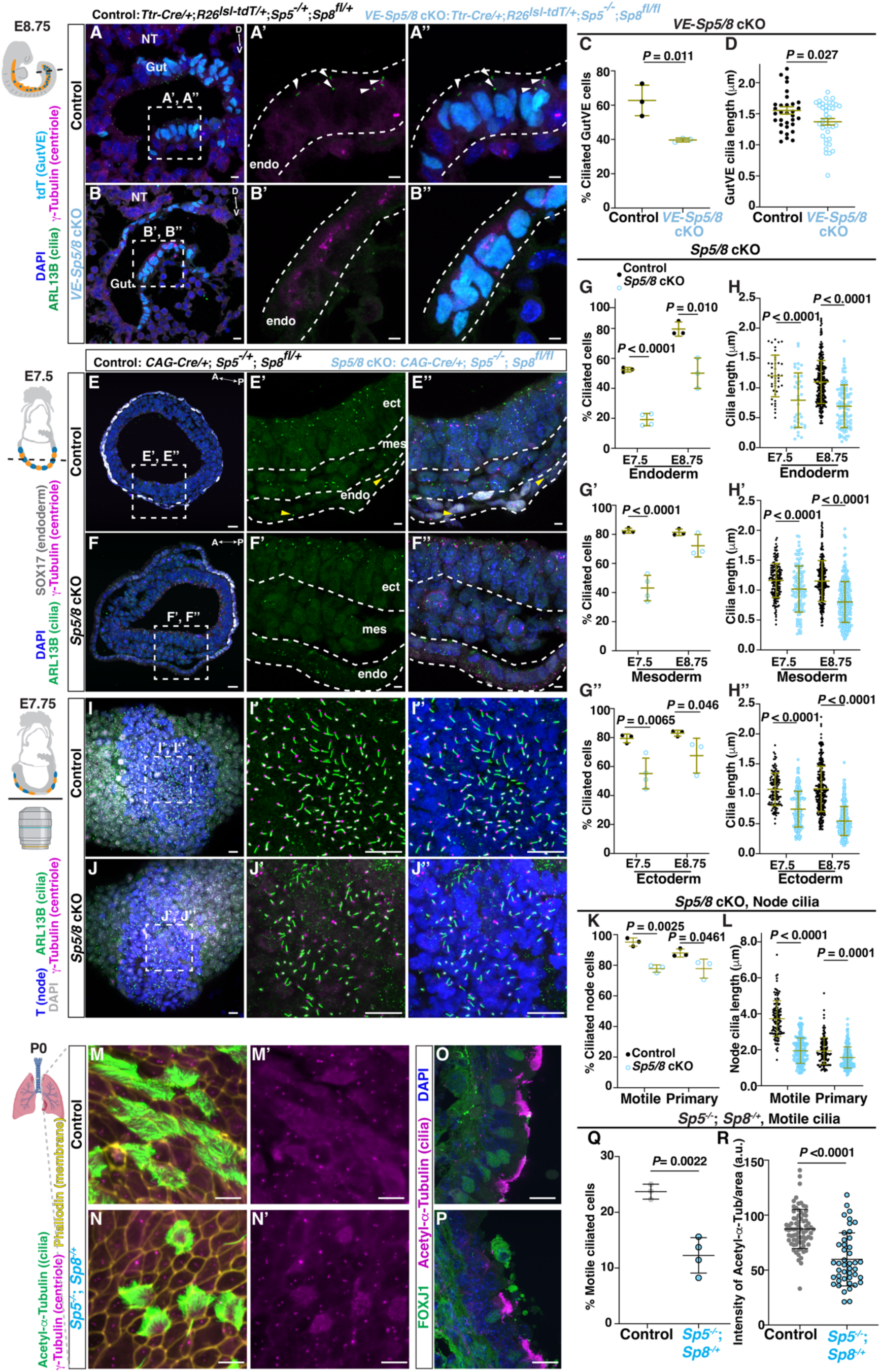
SP5/8 are required for primary cilia formation across tissues. (**A**, **B**) IF staining of transverse sections of Control and *VE*-*Sp5/8* cKO embryonic gut tube. Arrowhead, ciliated GutVE; NT, neural tube; D, dorsal; V, ventral. (**C**, **D**) Quantification of the percentage of ciliated cells and cilia length (D). (**E**, **F**) IF staining of transverse sections of Control and *Sp5/8* cKOs. Ect, ectoderm; mes, mesoderm; end, endoderm. Note, in the *Sp5/8* cKOs endoderm, ARL13B signal in cytoplasm is unspecific. (**G**, **H**) Quantification of the percentage of ciliated cells (G-G’’) and cilia length (H-H’’) in three cell types. n≥3 animals/genotype. (**I**, **J**) Maximum intensity projection ventral view of wholemount IF staining of Control (I) and *Sp5/8* cKO (J) embryos in the node. (**K**, **L**) Quantification of percentage node ciliated cells (K) and cilia length (L) based on their position and *Brachyury* (T) expression (N=3 embryos/genotype). Motile and primary cilia are distinguished based on their localization at the center of the node (node pitch) or peripheral to it. (**M**, **N**) En face imaging of distal part of trachea in control and *Sp5^-/-^*; *Sp8^-/+^* P0 neonates stained for cilia and membrane markers. Schematic created from Biorender.com. (**O**, **P**) IF staining of sagittal sections of trachea in control and *Sp5^-/-^*; *Sp8^-/+^* animals at P0. (**Q**, **R**) Quantification of the percentage of motile ciliated cells (Q), and intensity of Acetyl-α-Tubulin fluorescence (R). Scale bar, 20μm (A, B, E, F, I, J); 20μm (M-P); 5μm insets. The schematics of the embryos beside (A, E, I) were modified from (*21*). Statistical analyses: unpaired *t*-test (C, D, Q, R), one-way ANOVA (G, H, K, L).

We next employed *CAG-Cre* to globally delete *Sp8* in the *Sp5* null mutant background (*Sp5/8* cKO). At E10.5 *Sp5/8* cKOs displayed the expected truncation of the anterior head and posterior trunk and exencephaly seen in *Sp*8 null mutants (*34*), but the phenotype was more extreme in double mutants (fig. S8C). At E7.5, there was an approximately two-fold reduction in the percentage of ciliated cells in all three germ layers (endoderm, mesoderm, and ectoderm) of *Sp5/8* cKOs compared to controls, with the strongest defect in endoderm cells marked by SOX17 (GutVE and DE) (Fig. 4E-G). At E8.75, the percentage of cells with cilia continued to be reduced in *Sp5/8* cKOs, especially in the gut endoderm (Fig. 4G). Cilia length was also significantly decreased in *Sp5/8* cKOs in all three cell types, possibly more severely at E8.75 than E7.5 (Fig. 4H, fig. S8D, E). Importantly, the cilia defect was not due to lack of intercalation of VE and DE in the *Sp5/8* cKOs since intercalation was normal (fig. S8H-K). Furthermore, by analyzing single and homozygous/heterozygous double mutant combinations we found that *Sp5* and *Sp8* both contribute to cilia formation at E8.5 across the three cell types (fig. S8L, M). These results demonstrate a global requirement of SP5/8 in promoting primary cilia formation and length control.

Primary cilia are essential for Sonic hedgehog (SHH) signaling and act through GLI TFs to ventralize the neural tube (*40*). As expected for a cilia mutant, we found a dorsalization of the neural tube in E10.5 *Sp5/8* DKO mutants with a ventral expansion of PAX6+ dorsal progenitors into, and replacing, much of the OLIG2+ motor neuron domain (fig. S9). There was also a major reduction of the SHH target gene *Gli1*.

### *Sp5/8* promote motile cilia formation and function

In the early embryo, the node cells are a critical organizing center for gastrulation, and have a single specialized motile primary cilium that is longer than typical primary cilia and generates fluid flow necessary to generate organ asymmetry (*2*). We detected SP8 protein in the E7.75 node (fig. S10A, B) and *Sp5/8* cKO mutants exhibited a significant reduction in the percentage of node cells with motile cilia compared to controls (77.8% vs 95.2%, respectively) and a 50% reduction in node cilia length (1.9µm in mutants vs 3.7µm in controls) (Fig. 4I-L). Primary cilia number in the peripheral node region of *Sp5/8* cKO mutants was also reduced (77.8% in mutants vs 88.0%), with a shorter length (1.6µm in mutants vs 1.9µm in controls). Consistent with a defect in node cilia, *Sp5/8* double null mutants had altered left-right patterning, as shown by bilateral expression of *Nodal* and *Lefty1/2* at E8.25 (6 somites) (fig. S10C, 10D, E8.25) and randomization of heart looping at E9.5 (40% situs inversus/ambiguous) and lung morphology at E14.0 (57% situs inversus/ambiguous) compared to *Sp5^-/-^*; *Sp8^-/+^*controls (fig. S10E-G).

Given that the motile cilia genes *Foxj1* and *Rfx2/3/7* have binding sites for SP5/8, we next asked whether motile cilia in the lung endoderm and ependymal cells of the brain require *Sp5/8* for their normal formation. Although *Sp8* cKOs die at E14, we found that *Sp5^-/-^*; *Sp8^-/+^* animals die shortly after birth and have spina bifida and limb defects (fig. S11A). Most *Sp5^-/-^*; *Sp8^-/+^* animals also had hydrocephalus (fig. S11B). In the distal trachea at P0, we observed a reduction in the number of FOXJ1+ nuclei corresponding to a significant reduction in the percentage of cells with motile cilia and within those cells the intensity of acetylated α tubulin was reduced indicating less and shorter cilia per cell (Fig. 4M-R). The number of centrioles also appeared reduced. Similarly, in the brain we found a significant reduction in the percentage of FOXJ1+ ependymal cells lining the third ventricle, corresponding to a reduction in cilia (ARL13B staining) and centrioles per cell (fig. S11C-E). These findings demonstrate that SP5/8 play a crucial role in promoting motile cilia formation in addition to primary cilia.

### SP8 is sufficient to induce primary cilia in unciliated cells

We next tested whether SP8 is sufficient to induce cilia formation in YsVE cells by utilizing a Dox-inducible *Sp8* gain-of-function (GOF) mouse allele combined with *Ttr-Cre* (*24*) and *R26^lsl-rtTA-Ires-EGFP^* (*41*) alleles and administering Dox daily during E5.5 to 7.5 (*VE-Sp8* GOF; fig. S12A). In E8.75 *VE-Sp8* GOF embryos SP8 protein was detected in the YsVE (fig. S12B, C) and strikingly cilia were present on 12% of cells compared to none in controls (fig. S12D-G). Furthermore, cilia length was like nonmotile primary cilia (∼1.3µm).

A similar strategy was utilized using *CAG-Cre* to globally induce SP8 expression (*Sp8* GOF) (Fig. 5A) and again ∼15% of YsVE cells developed cilia. Interestingly, the percentage of YsMes with cilia also increased (Fig. 5B-D). Likewise, in Dox-inducible *i-Sp8-3xFlag* ES cells the percentage of cells with cilia was increased nearly two-fold from 22% to 35.0% and the average length of cilia was slightly increased (1.3µm in +Dox vs 1.1µm in -Dox, *P*=0.025, fig. S13A-C).

**Fig. 5.**
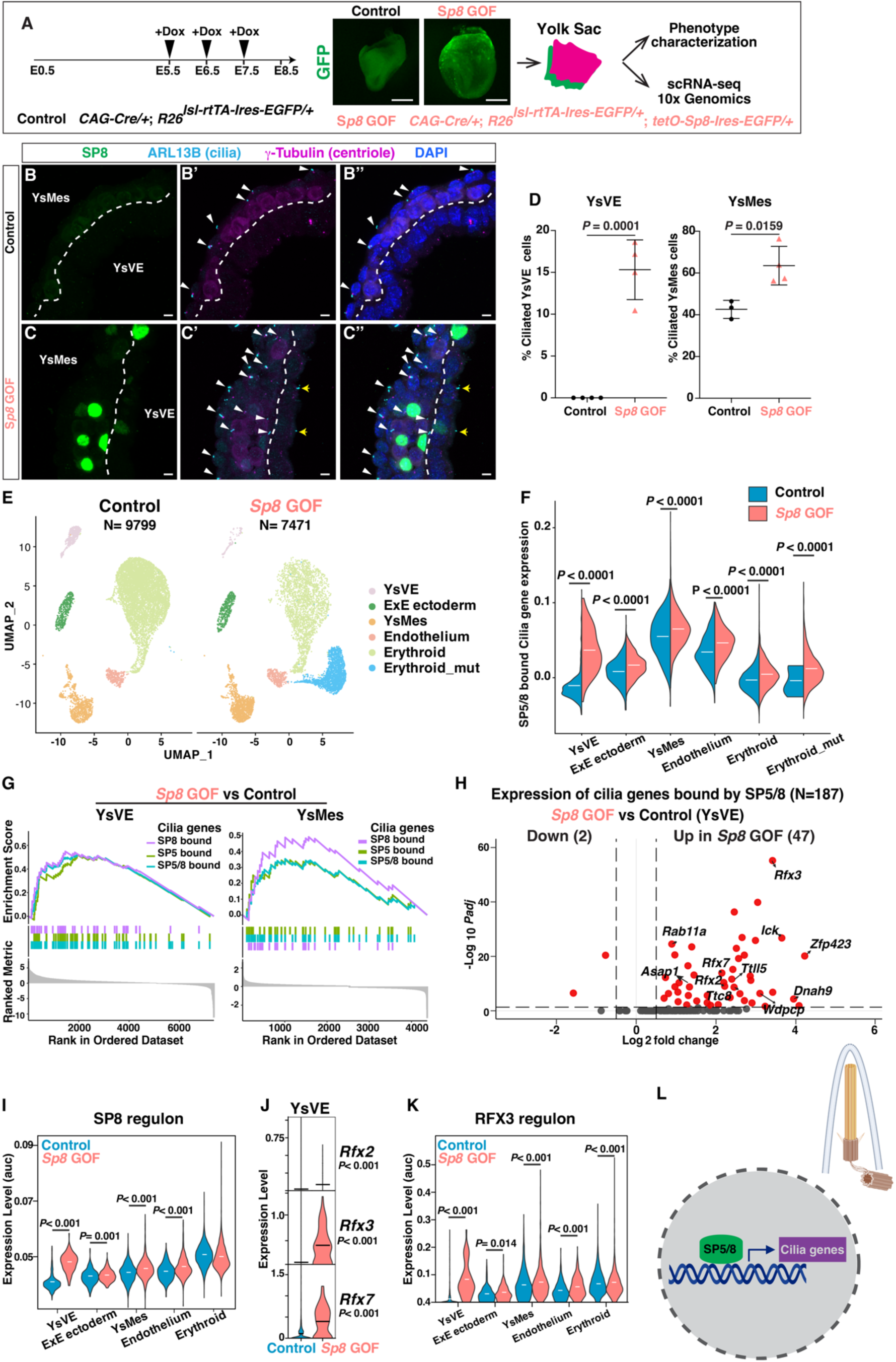
SP8 is sufficient to induce primary cilia in YsVE. (**A**) The strategy of global *Sp8* gain-of-function (GOF) (top) for phenotype characterization and scRNA-seq analysis in Control and *Sp8* GOF E8.75 embryos. (**B**, **C**) IF staining of transverse sections of yolk sac tissues from Control (B) and *Sp8* GOF (C) E8.5 embryos. White arrowheads, ciliated YsMes; yellow arrows, ciliated YsVE in *Sp8* GOF mutants. (**D**) Quantification of the percentage of ciliated YsVE and YsMes cells in Control and *Sp8* GOF yolk sacs (N≥3 tissues/genotype). (**E**) UMAP plots showing clustering of cell types of Control and *Sp8* GOF yolk sac tissues (N=3, 2 replicates). (**F**) Violin plots showing the gene set expression levels of SP5/8 bound cilia genes (N=187) across cell types in Control and *Sp8* GOF yolk sac cell types. (**G**) Gene set enrichment plots showing that SP5-bound, SP8-bound, or combined SP5/8-bound cilia genes are enriched in *Sp8* GOF group vs Control in YsVE and YsMes cell types. (**H**) Volcano plot showing the differential expression of SP5/8 bound cilia genes in YsVE cells, comparing *Sp8* GOF to Control (table S24). (**I**) Violin plots depicting the AUC (Area Under the Curve) score of SP8 regulon across different cell types in Control and *Sp8* GOF samples. (**J**) Violin plots showing log_10_ CPM (counts per million) expression levels of *Rfx2*, *Rfx3* and *Rfx7* in YsVE of Control and *Sp8* GOF samples. (**K**) Violin plot depicting the AUC score of RFX3 regulon across different cell types in Control and *Sp8* GOF samples. (**L**) Schematic showing SP5/8 bind and activate a set of primary cilia genes (N=130 for SP5, N=84 for SP8), and some motile cilia genes (N=9 for SP5, N=5 for SP8) in the nucleus to promote cilia formation (top right). Scale bars, 250 μm (A), 5μm (B, C). Statistical analysis: unpaired *t*-test (D, J), Wilcoxon rank-sum test (F, I, K).

SP8 mis-expression using the Dox system in unciliated extraembryonic endoderm (XEN) stem cells (*19*), a model for primitive endoderm (precursor of VE) also induced cilia, but only in 3.5% of cells with a cilia length of 1.8µm (fig. S13D-L). In addition, overexpression of *Rfx2* or *Rfx4* were not sufficient to induce cilia formation in XEN cells and ectopic RFX2/4 did not increase cilia formation in the presence of SP8 (fig. S13M). Thus, of these TFs that bind cilia genes and are increased in E8.75 ciliated cells, only SP8 induces cilia.

Mis-expression of SP8 did not appear to change the property of the YsVE since ciliated YsVE maintained expression of the YsVE-specific protein CDH1/E-cadherin (fig. S12D, E). To further investigate the consequences of ectopic cilia formation in *Sp8* GOF embryos, we performed scRNA-seq analysis of control (N=3) and *Sp8* GOF (N=2) yolk sacs at E8.5 (Fig. 5E, fig. S14A-D). Cluster analysis identified five major cell types in the control and one additional cluster in the mutants. Pseudotime trajectory analysis suggested the new mutant cluster is related to the erythroid lineage (fig. S14E). The overall expression of SP5/8 bound cilia genes was significantly enhanced across all cell types in *Sp8* GOF embryos (Fig. 5F). In particular, this gene set was highly enriched in YsVE and YsMes (Fig. 5G). Of the 187 cilia genes bound by SP5/8, 47 were significantly upregulated in *Sp8* GOF YsVE, including genes encoding IFT, BBS, RFXs, and other positive regulators of ciliogenesis (Fig. 5H, table S24). Indeed, the regulon activity of SP8 was greatly enhanced across all cell types but erythroid (Fig. 5I). *Rfx*2/*3*/7 were among the genes significantly increased in the YsVE, along with RFX3 target genes (Fig. 5J, K). We did not detect cell death in control or *Sp8* GOF conditions, thus SP8 does not induce cell death (fig. S15A). Lastly, although mutant YsVE cells form cilia, expression of *Gli2* and *Gli3* and the SHH target gene *Gli1* were very low and not induced in mutant YsVE, whereas they were increased in YsMes that has the machinery necessary for SHH-signaling. Thus, canonical SHH signaling is not induced in YsVE, although *Ihh* expression was increased (fig. S15B).

In summary, SP8 is sufficient to drive cilia formation in unciliated YsVE *in vivo* and XEN cells in culture and increase the percentage of ciliated cells, and this is accompanied by activation of SP5/8 target cilia genes (Fig. 5L).

### SP5/8 do not regulate cilia formation via the cell cycle

Cilia form during G1/GO and are disassembled before mitosis, and thus changes in cell cycle phases can alter cilia formation (*42–44*). In normal embryos where most cells are highly proliferative, we found a smaller percentage of ciliated cell types (DE and GutVE) are in G1 than unciliated YsVE at E6.5-8.75 (fig. S16A). Since SP5/8 have been reported to change cell cycle by promoting or inhibiting proliferation in a cell-type dependent manner (*45–47*), we asked whether loss or gain of SP5/8 alters the cell cycle in our systems. Analysis of *Sp8* GOF yolk sac sections did not detect a significant change in the percentage of YsVE cells in late G2-M (phospho-histone H3+) or S phase (EdU+ cells) when SP8 was over-expressed (fig. S16B-G). Cell cycle phase scoring of the scRNA-seq data also indicated the cell cycle phase distributions were not changed across cell types in *Sp8* GOF embryos compared to controls, except a possible increase in S phase in YsVE (fig. S16H). Flow analysis showed that the cell cycle distributions also were not altered in ES or XEN cells upon *Sp8* overexpression (fig. S17A-D). Furthermore, cell cycle arrest alone did not induce cilia formation in XEN cells or further enhance cilia formation in *Sp8* GOF XEN cells (fig. S17E). Finally, on the flip side, loss of *Sp5/8* in embryos did not change the percentage of cells in late G2-M or S phase in E8.5 embryonic neural ectoderm or reduce G1 in gastruloids (fig. S17F-J). Thus, SP5/8 do not regulate cilia formation through altering the cell cycle.

## Discussion

Previous work proposed that cilia formation is regulated in a lineage-specific manner, such that epiblast-derived cells have cilia and extra-embryonic lineage cells do not (*19*). While this is the case when the lineages are initially specified and segregated, we uncovered that when emVE intercalates with DE to become GutVE, the cells upregulate a set of cilia genes shared with other ciliated cells (YsMes and DE) and form primary cilia. This process is accompanied by upregulation of *Sp5/8* and we found many cilia genes have binding sites for SP5/8. Moreover, *Sp5/8* are required across embryonic cell types to increase the percentage of cells that form cilia and promote cilia length, for both primary and motile cilia. Finally, we demonstrate that a single TF, SP8, is sufficient to induce primary cilia in unciliated endoderm cells.

In addition to a reduction in the percentage of cells in the lung and brain that have motile in *Sp5^-/-^*; *Sp8^-/+^* neonates, the remaining cells have less cilia and centrioles per cell. Likewise, in *Sp5/8* cKO embryos, the length of the motile primary cilia remaining in the node are reduced two-fold but are longer than nonmotile primary cilia in other cell types. Consistent with these phenotypes, the mutants have hydrocephalus and randomization of left-right asymmetry of organs (lung and heart). Likely of relevance, we found that SP5/8 bind and SP8 can activate the expression of several motile cilia TFs (*Foxj1*, *Rfx2/3/7*). Since *Foxj1* and *Rfx3* null mutant mice have defects in node cilia length (*8, 15*), and *Rfx3* mutants have a 40% reduction in pancreas primary cilia and *Rfx4* mutants have shorter cilia in the embryonic brain (*13, 48*), the primary cilia defects observed in *Sp5/8* cKOs could in part be indirectly due to a decrease in expression of these TFs. Similarly, the defects in motile cilia length and number per cell in *Sp5/8* cKOs could be due to poor activation of *Foxj1* and *Rfx2/3/7* in the absence of SP5/8. However, our work argues that SP5/8 play a more prominent role than motile cilia TFs in primary cilia formation since in *Sp5/8* cKOs the percentage of ciliated cells and average length of cilia are significantly reduced across all cell types. Also, SP5/8 bind a larger number of primary cilia genes than FOXJ1 or RFXs (*12, 17, 49*).

Canonical WNT signaling was shown to enhance primary cilia formation in hTERT retinal pigment epithelium cells by stabilizing centriole satellite proteins (*50*). Our study shows that the WNT target gene *Sp5* positively regulates primary cilia formation in the mouse embryo and ES cells. We therefore suggest that WNT signaling acts, at least in part, at the transcriptional level by inducing *Sp5* gene expression and then SP5 induces expression of its target cilia genes to enhance cilia formation.

Given our findings that SP5/8 bind key TFs regulating motile cilia and motile cilia are reduced in *Sp5/8* mutants, SP5/8 are upstream in both the motile and primary cilia networks. However, although over-expression of SP8 in ES cells causes a slight increase in cilia length, the length of the extra cilia generated, or ectopic cilia in unciliated cells, is in line with primary cilia, and the cells have a single cilium. Thus, SP5/8 are not sufficient to induce motile cilia, at least in the time frame and systems we have analyzed. This is the case even when SP8 is co-expressed with RFX2 or RFX4 in XEN cells. Thus, SP5/8 are at the top of a transcriptional network activating primary and motile cilia genes but are only sufficient to induce primary cilia. *SP5/8* therefore should be considered as candidate genes mutated in human ciliopathies.

## Materials and Methods

### Mouse husbandry

The following mouse strains were used: *Afp-GFP* (*23*), *Ttr-Cre* (*24*), *R26^lsl-tdT/+^* (*25*), *Fgf8^CreER/+^*(*26*), *Strip1^fl/+^* (*27*), *Sp8^fl/fl^* (*33*), *Sp8^-/+^* were generated from *Sp8^fl/+^* mice*, tet-O-Sp8-Ires-EGFP* (*35*), *Sp5^lacZ/lacZ^*(*Sp5^-/-^*) (*39*), *R26^lsl-rtTA-Ires-EGFP^*(*41*), *CAG-Cre* (*51*), *Sox2-Cre* (*52*), *R26^mTmG/+^* (*53*). Animals were maintained on an outbred Swiss Webster background. For embryonic stage, noon of the day that a vaginal plug was detected was designated as E0.5. All animal experiments were performed per the protocols approved and guidelines provided by the Memorial Sloan Kettering Cancer Center’s Institutional Animal Care and Use Committee (IACUC). Animals were given access to food and water ad libitum and housed on a 12 hour light/dark cycle. Mice were randomly chosen for analysis with no exclusions and using both males and females. Because some genotypes have a phenotype, blinding was not always possible.

For inducible SP8 expression, Doxycycline (Sigma) was administrated by gavage (10mg/ml in water and 0.2ml per animal daily) to pregnant females from E5.5-7.5 to induce *Sp8* expression.

### Cell lines

*i-Sp8-3xFlag* ESCs were generated as previously reported (*35*). *Sp5/8* DKO ESCs were generated from R1 ESCs by Amaxa nucleofection (Lonza, VPH-1001) using CRISPR-cas9/gRNA ribonucleoprotein complex following manufacturer’s instructions. For targeting *Sp5*, two gRNAs: GGAGTAGCCCGGGGGCAACG and CGCTTCGCTTGTCCCGAGTG were used to delete a 3’ region of Exon-2 encoding a 3x Zn-finger domain and for *Sp8*, two guides: AGAGGAGTGGATCCCAACCT and GGACCCCCCTAACTGCGCCG were used to delete the coding region in Exon-3. The nucleofection of Cas9/gRNA complex and pMax-GFP cocktail solution was performed in 2,000,000 cells using Amaxa Nucleofector (Lonza). One day after nucleofection, GFP-positive cells were sorted using SONY SH800 (Sony), and 300,000 cells were plated onto gelatin-coated 150 mm dish for picking single colonies. Genomic deletion was confirmed by Sanger sequencing (Psomagen, Rockville, MD).

*i-Sp8-3xFlag* ESCs were maintained on 0.1% gelatin-coated dishes in ESC media that contained KnockOut DMEM (Gibco) supplemented with 15% fetal bovine serum (HyClone), 2mM L-glutamine (Gibco), 0.1mM β-mercaptoethanol (Gibco), 0.1 mM non-essential amino acids (Gibco), 1 mM sodium pyruvate (Gibco), 1% v/v penicillin and streptomycin (Gibco), 1,000 units/mL leukemia inhibitory factor (LIF, Millipore), 1μM PD0325901 and 3μM CHIR99021 (Stemgent). *Sp5/8* DKO and R1 ESCs were cultured in the ES medium described above except without 2i (PD0325901 and CHIR99021) as described (*37*).

*i-Sp8-3xFlag* EpiSCs were generated from *i-Sp8-3xFlag* ESCs following a reported protocol (*54*). Briefly, ESCs were disassociated using Accutase (0.5mM EDTA, Innovative Cell Technologies) and plated on fibronectin-coated plates (16.7µg/ml solution, Millipore) at a density of 17,500 cells/cm**^2^** on mitomycin-treated embryonic fibroblast feeder cells in N2B27 media supplemented with 12.5ng/ml heat-stable recombinant human β-FGF (PeproTech) and 20ng/ml activin A (PeproTech). EpiSC were cultured in the same medium on feeder cells for 2 passages before ChIP-seq. N2B27 media consisted of 50% DMEM-F12 (Gibco), 50% Neurobasal medium (Thermofisher), 0.5% N2 supplement (Thermofisher), 1% B27 supplement without vitamin A (Thermofisher), 2mM GlutaMAX (Gibco), 1% penicillin/streptomycin and 0.1µM β-mercaptoethanol.

Control and *VE*-*Sp8*-GOF XEN cell lines (N=3/genotype) were derived from E3.5 *Ttr-Cre/+*; *R26^lsl-rtTA-Ires-EGFP/+^* and *Ttr-Cre/+*; *R26^lsl-rtTA-Ires-EGFP/+^*; *tet-O-Sp8-Ires-EGFP/+* mouse blastocysts respectively using a protocol described previously (*55*). Briefly, mouse blastocysts were flushed from dissected uteri of pregnant females using embryo max M2 medium (Millipore). Blastocysts were then transferred using a mouth pipette onto a 0.1% gelatin-coated plate with mitomycin-treated mouse embryonic fibroblast feeder cells in ESC medium with 15% serum and LIF (Millipore). After 3 passages, feeder cells were removed and XEN cells were maintained on 0.1% gelatin-coated plastic dishes and cultured in RPMI1640 (pH 7.2) media (Gibco) supplemented with 15% fetal bovine serum, 2mM L-glutamine, 0.1mM β-mercaptoethanol, 0.1mM non-essential amino acids, 1mM sodium pyruvate and 1% v/v penicillin and streptomycin. The GOF of *Sp8* genotype was confirmed by GFP fluorescence after incubation with Doxycycline for 48 hours.

*VE*-*Sp8* GOF XEN cells were treated with cell cycle arrest drugs to block cell cycle progression using compounds listed in table S25.

Control and *Sp8* GOF XEN cells were transfected with lentivirus carrying Dox inducible expression modules for *Rfx2* (pCW57-RFP-P2A-MCS-Rfx2), *Rfx4* (TetO-FUW-Rfx4) or Empty vector (pCW57-RFP-P2A-MCS). Construct information is listed in table S25.

### Gastruloid sphere derivation

Gastruloids were derived from 300 R1 and *Sp5/8* DKO ES cells in 96 well ultra-low attachment dishes for 48 hours and treated with CHIR (3μM) from D2 to D2.5 (*38*). Gastruloid spheres were collected at Day 2.5 for single-cell RNA sequencing (paper in preparation (Chalamalasetty et al)). Immunostaining of cilia was performed using a previously published protocol (*37, 38*).

### Immunofluorescent staining, whole-mount embryo imaging and histology

For immunofluorescent (IF) staining of ESCs, cells were plated on gelatin-coated chambers (Ibidi), rinsed twice with PBS and fixed in 4% paraformaldehyde (PFA) for 10 min at room temperature (RT). After fixing, cells were washed twice with PBS and permeabilized with PBS containing 0.5% Triton X-100 (PBST) for 10 min at RT and blocked for 60 min with blocking buffer (0.1% PBST and 5% normal donkey serum).

For IF of embryo sections, embryos were fixed in 4% PFA at 4°C overnight, embedded in optimal cutting temperature compound (OCT), and cryosectioned at 12μm. Slides were air-dried for 10 min at RT and then washed in PBS for 10 min. Slides were permeabilized with 0.5% PBST for 10 min at RT and blocked with blocking buffer for 1 hour.

Primary antibodies were diluted in blocking buffer and placed on cells or slides overnight at 4℃. Cells or slides were washed three times in 0.1% PBST for 5 min and then incubated with secondary antibodies (Alexa Fluor conjugated secondary antibodies) diluted at 1:500 in blocking buffer at room temperature for 1 hour. Counterstaining was performed using DAPI (1:1000, Invitrogen). The cells or slides were washed three times in 0.1% PBST for 5 min each before cover slipping using Fluorogel mounting medium (Electron Microscopy Sciences). Primary and secondary antibodies and their related concentrations are listed in table S25.

For IF of whole-mount embryos, embryos were dissected and fixed with 4% PFA overnight at 4℃. After washing twice with PBS, embryos were permeabilized in 0.5% PBST for 10 min, washed, and blocked in blocking solution at RT for 1 hour, and incubated with the primary antibody diluted in blocking solution at 4°C overnight. Embryos were washed three times in 0.1% PBST at RT for 1 hour, incubated with secondary antibody at RT for 1 hour, washed three times in 0.1% PBST, and imaged in PBS on a glass-bottomed MatTek dish (MatTek). Chromogenic whole-mount in situ hybridization of E9.5 embryos and β-galactosidase staining of E14 lungs were performed as described previously (*36*).

For Hematoxylin and eosin staining, coronal sections of mouse brain were stained with Hematoxylin and Eosin (Thermo Fisher Scientific) (H&E) according to the protocol from the manufacturer. Images were collected on a Nanozoomer 2.0 HT slide scanner.

### Whole Mount In situ Hybridization

Split initiator probes (V3.0) for mouse *T*/*Bra*, *Nodal* and *Lefty1/2* were synthesized by Molecular Instruments, Inc. HCR-RNA Fluorescent In situ Hybridization was performed as described (*56*). Briefly, E8.25 embryos were harvested in PBS, fixed in 4% PFA overnight at 4°C and dehydrated. For HCR, embryos were rehydrated, permeabilized with 10mg/ml proteinase K for 15 min at room temperature and post-fixed in 4% PFA for 20min. Embryos were pre-incubated in probe hybridization buffer followed by an overnight incubation in probe solution at 37^0^C. Embryos were washed the next morning and pre-incubated in amplification buffer. For amplification, embryos were incubated in 60 pmol of each hairpin per 0.5ml of amplification buffer for 16 hour at RT in the dark (*36*). Embryos were washed in 5x SSCT buffer and stained in 0.5mg/ml of DAPI solution in 5x SSCT buffer overnight at RT. The stained embryos were embedded on glass bottom dishes (MatTEK) and oriented in 1% ultra-low-melt agarose. Embryos were cleared using Ce3D++ solution at RT for at least 2-3 days in the dark.

RNA in situ hybridization of *Gli1* mRNA in section was performed as described using antisense RNA probes for *Gli1* (*57*). The templates for *Gli1* were generated by PCR using primers containing T3 polymerase promoters from postnatal cerebellum cDNA.

### TUNEL (terminal deoxynucleotidyl transferase dUTP nick end labeling) assay

Slides were first permeabilized with 0.5% Triton X-100, and pre-incubated with Tdt buffer (30 mM Tris-HCl, 140 mM sodium cacodylate and 1 mM CoCl_2_) for 15 min at room temperature. Slides were then incubated in a TUNEL reaction solution [containing terminal transferase (Roche, 3333574001) and Biotin-16-dUTP (Sigma-Aldrich, 11093070910)] for 1 hour at 37°C. Following the TUNEL reaction, slides were incubated in a Streptavidin Alexa Fluor 647 conjugate (Invitrogen, S-32357) for 1 hour. Slides were then washed in PBS twice for 5 min each followed by a 10 min incubation in DAPI. Prior to coverslipping using Fluorogel mounting medium (Electron Microscopy Science), slides were washed with PBS twice for 5 min each.

### EdU labeling and detection

For cultured cells, 10 μM EdU (5-ethynyl-2’-deoxyuridine) was added to the culture medium of ESCs and XEN cells for 1hour before collection. Cells were then disassociated with 0.05% Trypsin, fixed with 4% PFA for 10 min, followed by labeling with Click-IT kit using Sulfo-Azide-Cy5 (Lumiprobe cat: A3330) and DAPI. The cells were then applied to flow cytometry analysis (Fortessas, BD). FlowJo software was used to visualize and analyze the intensity of DAPI and EdU-Cy5 fluorescence.

To quantify cell proliferation in the neural tube of embryos, EdU (Invitrogen) was injected intraperitoneally into dams at 100 mg/g 1 hour before euthanasia. A Click-it EdU kit using Sulfo-Azide-Cy5 (Lumiprobe Corporation A3330) was used per the manufacturer’s protocol to stain sections.

### Transmission electron microscopy

Transmission electron microscope was performed as described previously (*58*). Briefly, yolk sac tissues were collected from E8.75 mouse embryos, fixed in 2% PFA, 2.5% glutaraldehyde, and 2 mM calcium chloride in 0.075 M sodium cacodylate buffer, pH 7.4. Yolk sac tissue was then treated with 0.1% tannic acid and washed in sodium cacodylate buffer, postfixed with 1% osmium tetroxide for 1 hour, and stained en bloc with uranyl acetate for 30 min. Tissue was then dehydrated in a graded series of ethanol, washed with acetone, and embedded in resin (Eponate12; Electron Microscopy Sciences). After polymerization at 60°C for 48 hours, ultrathin serial sections were cut, poststained with 2% uranyl acetate and 1% lead citrate, and imaged in a TEM (100CX; Jeol) with a digital imaging system at room temperature (XR41-C; AMT). Images were acquired at 80 kV. To improve clarity, some images were adjusted in Photoshop (Adobe Systems). In every case, all pixels in the image were adjusted uniformly, and all panels within a figure were adjusted with identical settings.

### Imaging

Wholemount embryos and sections were digitally imaged on a Nikon A1R HD25 laser scanning confocal microscope and deconvolved using Denoise AI tool on NIS-Elements. Sections and images for quantification were imaged on an inverted DeltaVision Image Restoration Microscope with a deconvolution function. For whole mount in situ imaging, acquired optical sections (3.3µm) were analyzed with Imaris software (Bitplane, Inc).

### Reverse Transcription Quantitative PCR

RNA was extracted from cells using the Trizol standard method (Invitrogen). Coding DNA was synthesized from 1-2.5μg total RNA of biological triplicates (per each condition) by reverse transcription using oligo(d)T and random primers with SuperScript IV VILO (Thermofisher). The cDNA samples were diluted at 1:10, and 2.5μl of the diluted cDNA was used for reverse transcription quantitative PCR (RT-qPCR) of the candidate genes with SYBR Green Master Mix (Qiagen) on a StepOnePlus Real-Time PCR System (Life Technologies). Relative expression was calculated using the 2^-ΔΔCt^ method. The relative mRNA level was normalized to the reference gene *Gapdh* for data normalization. (Sequence of primers listed in table S25).

### Assay for Transposase-Accessible Chromatin (ATAC) and sequencing

ATAC-seq was performed as previously described (*59*), with minor modifications. The yolk sac and gut tube were dissected from E8.75 (12 somite) *Afp-GFP/+* embryos. Yolk sacs or gut tubes were washed in ice-cold DMEM/F12 and incubated for 45 min and 20 min, respectively, at 37°C in Accutase/0.25% Trypsin (Gibco) (1:2) to dissociate cells into single cells. Single-cell suspensions were incubated with DAPI to label dead cells for 5 min on ice and filtered through cell strainers prior to performing fluorescence-activated cell sorting (Aria-51, BD) to obtain high viability YsVE (GFP^+^) and YsMes (GFP^-^) cells from yolk sac samples and GutVE (GFP^+^) from gut tube samples.

For library preparation from the sorted cells described above was prepared according to the Omni-ATAC protocol. Briefly, 10,000-30,000 cells per sample were lysed using 0.1% NP-40, 0.1% Tween, and 0.01% Digitonin (Promega) to yield nuclei. The resulting chromatin was fragmented and tagmented using Tn5 transposase (Illumina). DNA was purified using a Qiagen MinElute Reaction Cleanup Kit (Qiagen) and amplified using NEBNext® High-Fidelity 2X PCR Master Mix (NEB). The number of cycles was estimated by qPCR. DNA tagmentation efficacy was evaluated with a Bioanalyzer 2100 (Agilent Technologies) and the DNA amounts were calculated with Qubit. Libraries were prepared using universal forward and reverse primers from Ad2.1-Ad2.24. The final libraries were purified using a single left-handed bead purification with AMPure beads (Beckman Coulter). The resulting DNA libraries were sequenced using the NextSeq550 system (Illumina), and about 25 million reads were obtained per sample by the MSKCC IGO core facility.

### Chromatin immunoprecipitation (ChIP) and sequencing

*i-Sp8-3xFlag* EpiSCs were treated with or without Doxycycline (1μg/ml) for 24 hours, and fixed in 1% formaldehyde for 10 min, after which the reaction was quenched by the addition of glycine to the final concentration of 0.125M. Fixed cells were washed twice with PBS, snap frozen and store at −80°C. Frozen cell pellets were sent to MSKCC Epigenetics Research Innovation Lab for processing. Cells were resuspended in SDS buffer (100mM NaCl, 50mM Tris-HCl pH 8.0, 5mM EDTA, 0.5% SDS, 1x protease inhibitor cocktail from Roche). The resulting nuclei were spun down, resuspended in immunoprecipitation buffer at 1 mL per 0.5 million cells (SDS buffer and Triton Dilution buffer (100mM NaCl, 100mM Tris-HCl pH 8.0, 5mM EDTA, 5% Triton X-100) mixed in 2:1 ratio with the addition of 1xprotease inhibitor cocktail (Millipore Sigma, #11836170001) and processed on a Covaris E220 Focused-ultrasonicator to achieve an average fragment length of 200-300 bps with the following parameters: PIP=140, Duty Factor=5, CBP/Burst per sec=200, Time=1800s. Chromatin concentrations were estimated using the Pierce™ BCA Protein Assay Kit (ThermoFisher Scientific) according to the manufacturer’s instructions. Chromatin equal to 25M cells was used per ChIP reaction. The immunoprecipitation reactions were set up in 500uL of the immunoprecipitation buffer in Protein LoBind tubes (Eppendorf) and pre-cleared with 50uL of Protein G Dynabeads (ThermoFisher Scientific) for 2 hours at 4°C. After pre-clearing, the samples were transferred into new Protein LoBind tubes and incubated overnight at 4°C with 5ug of Flag antibody (Millipore Sigma) and H3K4me3 antibody (Epicypher). The next day, 50μL of BSA-blocked Protein G Dynabeads were added to the reactions and incubated for 2 hours at 4°C. The beads were then washed two times with low-salt washing buffer (150mM NaCl, 1% Triton X-100, 0.1% SDS, 2mM EDTA, 20mM Tris-HCl pH8.0), two times with high-salt washing buffer (500mM NaCl, 1% Triton X-100, 0.1% SDS, 2mM EDTA, 20mM Tris-HCl pH8.0), two times with LiCL wash buffer (250mM LiCl, 10mM Tris-HCl pH8.0, 1mM EDTA, 1% Na-Deoxycholate, 1% IGEPAL CA-630) and one time with TE buffer (10mM Tris-HCl pH8.0, 1mM EDTA). The samples were then reverse-crosslinked overnight in the elution buffer (1% SDS, 0.1M NaHCO_3_) and purified using the ChIP DNA Clean & Concentrator kit (Zymo Research, #D5205) following the manufacturer’s instructions. After quantification of the recovered DNA fragments, libraries were prepared using the ThruPLEX®DNA-Seq kit (R400676, Takara) following the manufacturer’s instructions, purified with SPRIselect magnetic beads (B23318, Beckman Coulter), and quantified using a Qubit Flex fluorometer (ThermoFisher Scientific) and profiled with a TapeStation (Agilent). The libraries were sent to the MSKCC Integrated Genomics Operation core facility for sequencing on an Illumina NovaSeq 6000 (aiming for 30-40 million 100bp paired-end reads per library).

### ChIP and ATAC-seq data analysis

ChIP and ATAC sequencing reads were trimmed and filtered for quality and adapter content using version 0.4.5 of TrimGalore (https://www.bioinformatics.babraham.ac.uk/projects/trim_galore), with a quality setting of 15, and running version 1.15 of cutadapt and version 0.11.5 of FastQC. Reads were aligned to genome assembly mm10 using Bowtie2 (v2.3.4.1, http://bowtie-bio.sourceforge.net/bowtie2/index.shtml) and were deduplicated using MarkDuplicates (v2.16.0) of Picard Tools. To ascertain regions of chromatin accessibility, MACS2 (https://github.com/taoliu/MACS) was run with a p-value setting of 0.001 against a matched input sample. The bedtools suite (http://bedtools.readthedocs.io) was used to create normalized read density profiles by extending the 3’ end of the aligned fragments 200 bp for ChIP and 0 bp for ATAC and then normalizing to 10M uniquely mapped reads. A global peak atlas was created by first removing blacklisted regions (blacklists/mm10mouse/mm10.blacklist.bed.gz), then taking 500 bp windows around peak summits for ATAC and the entire peak region for ChIP and counting reads with version 1.6.1 of featureCounts (http://subread.sourceforge.net). DESeq2 was used to normalize read density (median ratio method) and to calculate differential enrichment for all pairwise contrasts. Peaks were annotated to genomic elements such as transcription start sites (TSS), transcription termination sites (TTS), exons, and introns using GENCODE version M17. Peak intersections were calculated using bedtools v2.29.1 and intersectBed with 1 bp overlap. Motif signatures were obtained using Homer v4.5 (http://homer.ucsd.edu). Composite and tornado plots were created using deepTools v3.3.0 by running computeMatrix and plotHeatmap on normalized bigwigs with average signal sampled in 25 bp windows and flanking region defined by the surrounding 2 kb. Peaks (regions of enrichment) were visualized using IGV (v2.16.0).

### Single cell RNA-sequencing (scRNA-seq)

For the normal E8.75 embryos sample preparation, single cells were obtained as previously reported (*15*). Briefly, yolk sacs from three E8.75 embryos (12 somites) were dissected in DMEM/F12 with 5% FBS. Cells were dissociated with Trysin/Accutase for 15 min at 37°C. Cell clumps were triturated into single cells by mouth-pipetting using pulled (Sutter Instruments) 75mm glass capillaries. To remove debris, single-cell suspensions were filtered through FlowMI cell strainers (40μm, Sigma-Aldrich). Single cells were then spun at 450g for 4 min at RT and resuspended in PBS+0.04% BSA. Cell numbers and viability were determined using 0.2% (w/v) Trypan Blue on a hemocytometer. Single-cell suspensions of biological repeats were labeled with a Cell Multiplexing Oligo (10x Genomics 3’ CellPlex Kit) and then pooled together prior to loading onto a 10x Genomics chip (Chromium Next GEM Chip G). Following the manufacturer’s instructions, single-cell libraries were prepared using 3’ CellPlex Kit. After PicoGreen quantification and quality control by Agilent TapeStation, final libraries were sequenced on Illumina NovaSeq S4 platform (R1-28 cycles, i7-8 cycles, R2-90 cycles). The cell-gene count matrix was constructed using the Sequence Quality Control (SEQC) package (*60*).

For *Sp8* GOF scRNA-seq, single cells were obtained from E8.5 Control and *Sp8* GOF embryos as described above. Yolk sac tissues from 2 embryos per sample (N=2 per condition) were dissociated and filtered as described above. Single-cell suspensions were incubated with DAPI to label dead cells for 5 min on ice and filtered through cell strainers, followed by fluorescence-activated cell sorting (Aria-51, BD) to obtain single and high-viability cell suspensions. 9K-15K single-cell suspensions were loaded onto a 10x Genomics chip (Chromium Next GEM Chip G).

Following the manufacturer’s instructions, single-cell libraries were prepared using 10X Genomics’ Chromium Next GEM Single Cell 3’ Reagent Kits, v3.1. After PicoGreen quantification and quality control by Agilent TapeStation, final libraries were sequenced on Illumina NovaSeq X platform. The cell-gene count matrix was constructed using the Sequence Quality Control (SEQC) package (*60*).

### scRNA-seq data analysis

For E8.75 yolk sac scRNA-seq data, the Cell Ranger Single Cell software suite (10x Genomics) was used to demultiplex samples, align reads, and generate feature-barcode matrices. The reference genome used was the Genome Reference Consortium Mouse Build 38 (GRCm38, Gencode annotation mm10). Raw reads were processed using the Cell Ranger count program using default parameters.

For E8.5 *Sp8* GOF yolk sac scRNA-seq data, the Cell Ranger Single Cell software suite (10x Genomics) was used to align reads and generate feature-barcode matrices. The reference genome used was the Genome Reference Consortium Mouse Build 38 (GRCm38, Gencode annotation mm10). Raw reads were processed using the Cell Ranger count program using default parameters.

For both of these scRNA-seq studies, Seurat v4.3.0 package was used to generate a UMI (unique molecular identifier) count matrix from the Cell Ranger output (*54*). Genes expressed in less than 10 cells were removed for further analyses. Cells with larger than 500 UMIs, 250 genes, 0.20 mitoRatio, or log10GenesPerUMI ≤ 0.75 were considered low quality/outliers and discarded from the datasets. For the E7.5 yolk sac analysis, normalization was performed on individual samples using the logTransform function. For the E8. 5 *Sp8* GOF yolk sac scRNA-seq data normalization was performed on individual samples using the NormalizeData function with default parameters. The normalized data was scaled by ScaleData function with mitochondrial genes percentage regressed out. Principal component analysis (PCA) was performed on the scaled data by runPCA. Samples were then integrated using IntegrateLayers function with the Harmony Integration method.

Samples were then integrated using the Harmony Integration functions. For all analyses, the first 15 dimensions were used for the FindNeighbors function, and clusters were identified using the FindClusters function with a resolution of 0.1. Data were projected into the 2D space using the FindUMAP or RunUMAP function with 15 dimensions. Cluster markers and further differential gene expression analyses were all performed using normalized counts (NormalizeData) in the RNA assay. Cluster markers were identified using the FindAllMarkers and comparing markers generated to existing literature. To refine clustering further, the SubsetData function was used to create a new Seurat object, and the above clustering was reiterated.

For the gastruloid sRNA-seq analysis, we used STARSolo (STAR v. 2.7.9a) to map control and *Sp5/8* DKO libraries to the same reference genome (*63*). Raw count matrices were processed with CellBender v. 0.3.2 using default parameters (*64*), then used to create individual Seurat v. 5.1.0 objects. For each object, we removed genes expressed in fewer than three cells and cells with fewer than 200 genes, and excluded cells beyond three median absolute deviations for mitochondrial transcript percentage, feature count, or UMI count. We then used DoubletFinder v. 2.0.4 to remove the top 7.5 % most likely doublets (*65*). Filtered objects were merged, log-normalized (NormalizeData, scale = 10 000), variable features identified (FindVariableFeatures, n = 2 000), and data scaled (ScaleData, regressing out mitochondrial RNA and cell-cycle scores). We ran PCA on the 2 000 variable features (RunPCA), integrated samples via IntegrateLayers with Harmony, and identified differentially expressed genes between control and *Sp5/8* DKO using FindMarkers.

For publicly available scRNA-seq data, the endoderm scRNA-seq raw data were downloaded from GSE123046 (*21*), and the Seurat package was used to generate a Seurat object following the standard workflow described above. DE (E7.5 and E8.75 DE derived gut endoderm cells), emVE + GutVE (E5.5 to E7.5, and E8.75 GutVE derived gut endoderm cells), and exVE + YsVE (E5.5 to E8.75) were further subsetted and re-clustered for downstream data visualization and differential gene expression analysis.

Differential gene expression analyses between GutVE, DE, and YsMes compared to YsVE were performed using the Libra algorithm with the edgeR-LRT pseudobulk method (*61*). Genes with an adjusted *P* value (*Padj*) < 0.05 were considered significantly up or downregulated. Results were visualized by Violin plot and Umap plot using Seurat package, and Volcano plot using the EnhancedVolcano package on Bioconductor. Gene ontology analyses were performed separately on up and downregulated genes using the enrichGO, clusterProfiler packages on Bioconductor (*62*). To build a pseudotime trajectory on UMAP, Seurat object was converted to monocle 3 (1.3.7) object by as.cell_data_set function(*66, 67*). Cells were then clustered by cluster_cells function. The trajectory each partition was constructed by learn_graph followed by order_cells function. Regulon activity was computed by SCENIC (1.3.1) R package as previously described (*68*).

Human CS7 scRNA-seq data was downloaded from ArrayExpress (E-MTAB-9388) (*30*), and the Seurat package was used to generate a Seurat object following the standard workflow. YsMes, Gut endoderm and YsE cell types were further subsetted using Subset function, and plot gene expression using Doheatmap function of Seurat package.

### Statistics

All statistical analyses of quantified data were performed using Prism software (GraphPad v10.2.3). The statistical analysis used for each figure is described in the figure legends. Data (shown as scatter graphs) are presented as mean±standard deviation (S.D.). Student unpaired Two-tailed *t*-test was used to compare two conditions; one-way ANOVA using an FDR method (Benjamini and Hochberg) was used to compare two experimental groups; two-way ANOVA followed by Tukey’s multiple comparison test was used for group analysis. *P* values were indicated in the figure/legends. The statistical significance of the SP5/8 ChIP tracks overlap with the cilia gene open chromatin, and of ATAC-seq genes/peaks overlap with cilia gene list was calculated using a two-sided Fisher’s exact test. The statistical significance of gene/gene module expression of scRNA-seq data was calculated using Wilcoxon rank-sum test.

## Acknowledgments

We thank members of Joyner lab for helpful discussions and especially Anjana Krishnamurthy and Sumru Bayin for providing advice on the scRNA-seq experiments and Andrew Lee for advice on analyzing cilia in the brain. We also thank Junmin Pan (Tsinghua University) for insightful comments on the manuscript; Meng-Fu Bryan Tsou (MSKCC) for comments on the manuscript and sharing CEP162 antibody; Jennifer Zallen (MSKCC) for providing *R26^mTmG^*; *Strip1^fl/fl^* mice; Sonja Nowotschin for providing an Illustrator file from which to modify the schematics of embryos; Yosip Kelemen (Weill Cornell Medicine) for data processing for Regulon analysis; Bumsoo Kim and Joo-Hyeon Lee (MSKCC) for advice on trachea staining and data analysis. We are grateful to the following cores for their expertise: the Rockefeller University EM facility for TEM sample preparation and imaging, and the MSKCC Epigenetics Research Innovation Lab, Single-cell Analytics Innovation Lab (SAIL), Integrated Genomics Operation, Center for Comparative Medicine and Pathology, and Flow cytometry facility. This work utilized the computational resources of the NIH HPC Biowulf cluster (https://hpc.nih.gov). Y.L. and A.L.J. dedicate this manuscript to the memory of their late colleague Kathryn V. Anderson who provided continuous scientific inspiration and support. Kathryn also provided valuable mentorship to YL in the early stages of the work.

## Funding

National Institutes of Health grant R01GM126124 (ALJ, KVA)

National Institutes of Health grant R01HD035455-24 (ALJ, KVA, AKH)

National Institutes of Health grant CCSG, P30 CA08748 (ALJ, AKH, SV)

National Institutes of Health grant R01DK127821 (AKH)

Cycle for Survival GC-223397 (ALJ)

National Institutes of Health NCI, 1ZIABC010345 (TPY)

## Author contributions

Conceptualization: ALJ, YL

Methodology: ALJ, RK, XH, YL

Software: FPL, RK, XH, YL

Formal Analysis: YL, YP

Investigation: AKH, DS, RBC, ML, MWK, Y-YK, YL, ZL

Resources: ALJ, AKH, TPY

Visualization: ALJ, RK, RBC, XH, YL

Funding acquisition: AKH, ALJ, KVA, TPY

Supervision: AKH, ALJ, KVA, TPY

Project administration: ALJ, YL Writing – original draft: ALJ, YL

Writing – review & editing: ALJ, RBC, TPY, YL

## Competing interests

The authors declare no competing financial interests.

## Data and Materials Availability

Raw fastq sequencing files and output files of normal and *Sp8* GOF yolk sac scRNA-seq experiments are available at the GEO repository under the accession numbers GSE274844 and GSE293888. The scRNA-Seq data for the Gastruloid spheres are available at the GEO repository under the accession numbers GSE296732. Raw fastq sequencing files for ATAC-seq, ChIP-seq datasets and bigwig files for displaying read density tracks have been submitted to the GEO repository under the accession numbers GSE273919, GSE273920. All deposited data will be publicly available as of the date of publication. This paper does not report any original code; code resources and minor modifications are described in the Methods. Scripts for *Sp8* GOF yolk sac scRNA-seq data analysis are available at http://github.com/stevehxf/Cilia_Science2025.

Any materials not publicly available through GEO or GitHub are available by contacting ALJ or YL.

## Supplementary Materials

**Fig. S1.**
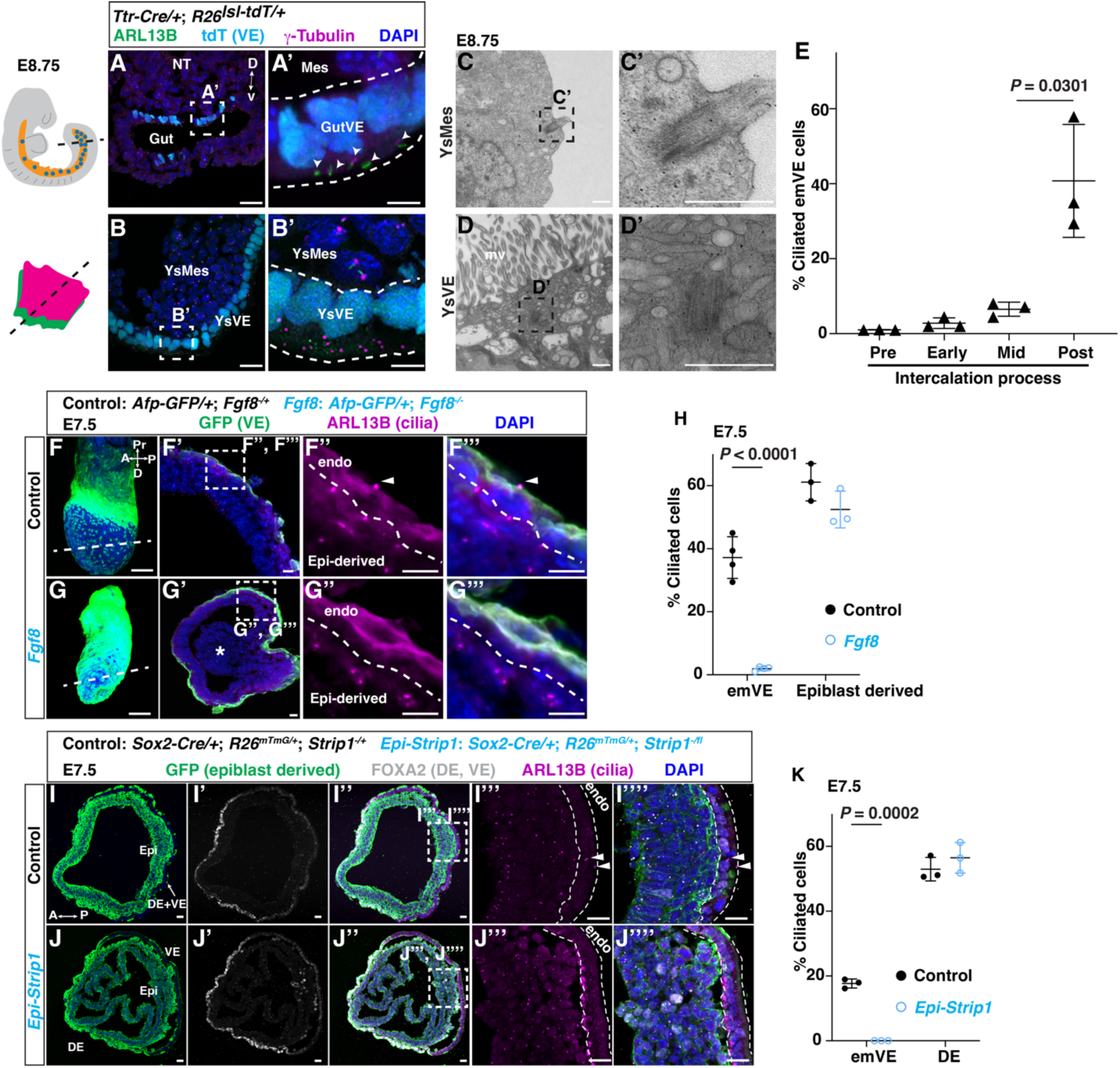
GutVE forms cilia during intercalation with DE. (**A**, **B**) IF staining of sections of gut tube (A) and yolk sac (B) from *Ttr-Cre/+*; *R26^lsl-tdT/+^* E8.75 embryos. (A’, B’) Higher magnification in (A, B). NT, neural tube. Arrowheads, ciliated GutVE. The schematic of the embryo beside (A) was modified from (*21*). (**C**, **D**) Transmission electron microscope images of yolk sac. (C’, D’) Dashed boxes, centriole/basal body at higher magnification. Scale bar, 500nm; mv, microvillus. (**E**) Quantification of the percentage of ciliated emVE cells based on the process of intercalation with DE, defined by embryo stage E6.5 (Pre), E7.0 (Early), E7.25 (Mid), and E7.75 (Post). (N=3 embryos/age). (**F**, **G**) Wholemount staining littermate Control and *Fgf8* null mutants. Asterisk, epiblast cells in the primitive streak that fail to undergo epithelial to mesenchymal transition. (**H**) Quantification of the percentage of ciliated cells in Control emVE or DE cells and in *Fgf8* null emVE or adjacent epiblast cells (N=3 embryos/genotype). (**I**, **J**) IF analysis of transverse sections of E7.5 Control and *Epi*-*Strip1* embryos. (**K**) Quantification of the percentage of ciliated cells in emVE (GFP^-^) and DE (GFP^+^, FOXA2^+^) of Control and *Epi*-*Strip1* embryos (N=3 embryos/genotype) in the posterior half of the embryo. A, anterior; P, posterior; Pr, proximal; D, distal; endo, endoderm; arrowheads, ciliated emVE. (I’’’, I’’’’, J’’’, J’’’’) higher magnification images of insets in (I’’, J’’). Scale bars, 500nm (C, D), 100μm (F, G), 20μm (A, B, F’, G’, I, J), 5μm in the insets. Statistical analysis: one-way ANOVA (E) or unpaired *t*-test (H, K).

**Fig. S2.**
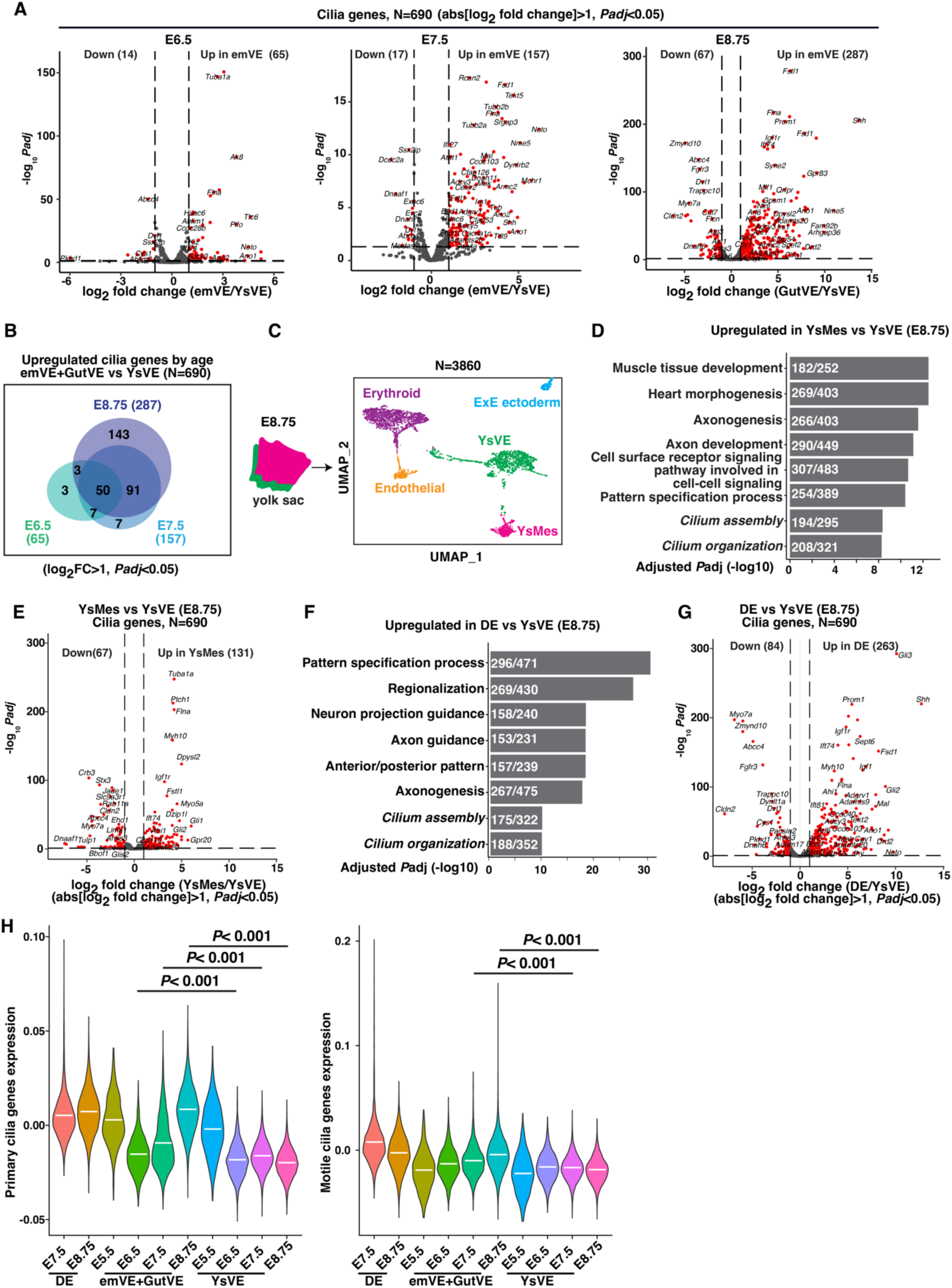
Cilia genes are upregulated in GutVE, DE, and YsMes compared to YsVE. (**A**) Volcano plot of scRNA-seq data showing the differentially expressed cilia genes (N=690) between emVE+GutVE and YsVE at three stages. (**B**) Venn diagram showing the overlap of upregulated cilia genes in emVE/GutVE compared to YsVE. (**C**) Schematic showing sample preparation for scRNA-seq (left) and UMAP plot showing the clusters (right), N=3 yolk sacs. (**D**) Bar plot showing top eight upregulated biological process terms and cilia-related terms in YsMes compared to YsVE at E8.75 (input gene cutoff was log_2_FC>0, *Padj*<0.05). (**E**) Volcano Plot of differential cilia gene expression in YsMes compared to YsVE at E8.75. (**F**) Bar plot showing top eight upregulated biological process terms and cilia-related terms in DE compared to YsVE at E8.75 (input gene cutoff was log_2_FC>0, *Padj*<0.05). (**G**) Volcano plot showing differentially expressed cilia genes in DE compared to YsVE at E8.75. **(H**) Violin plots showing the expression level of primary cilia genes and motile cilia genes in DE, GutVE, and YsVE at E5.5-E8.75 (gene listed in table S3). Statistical analysis: Wilcoxon rank-sum tests (H).

**Fig. S3.**
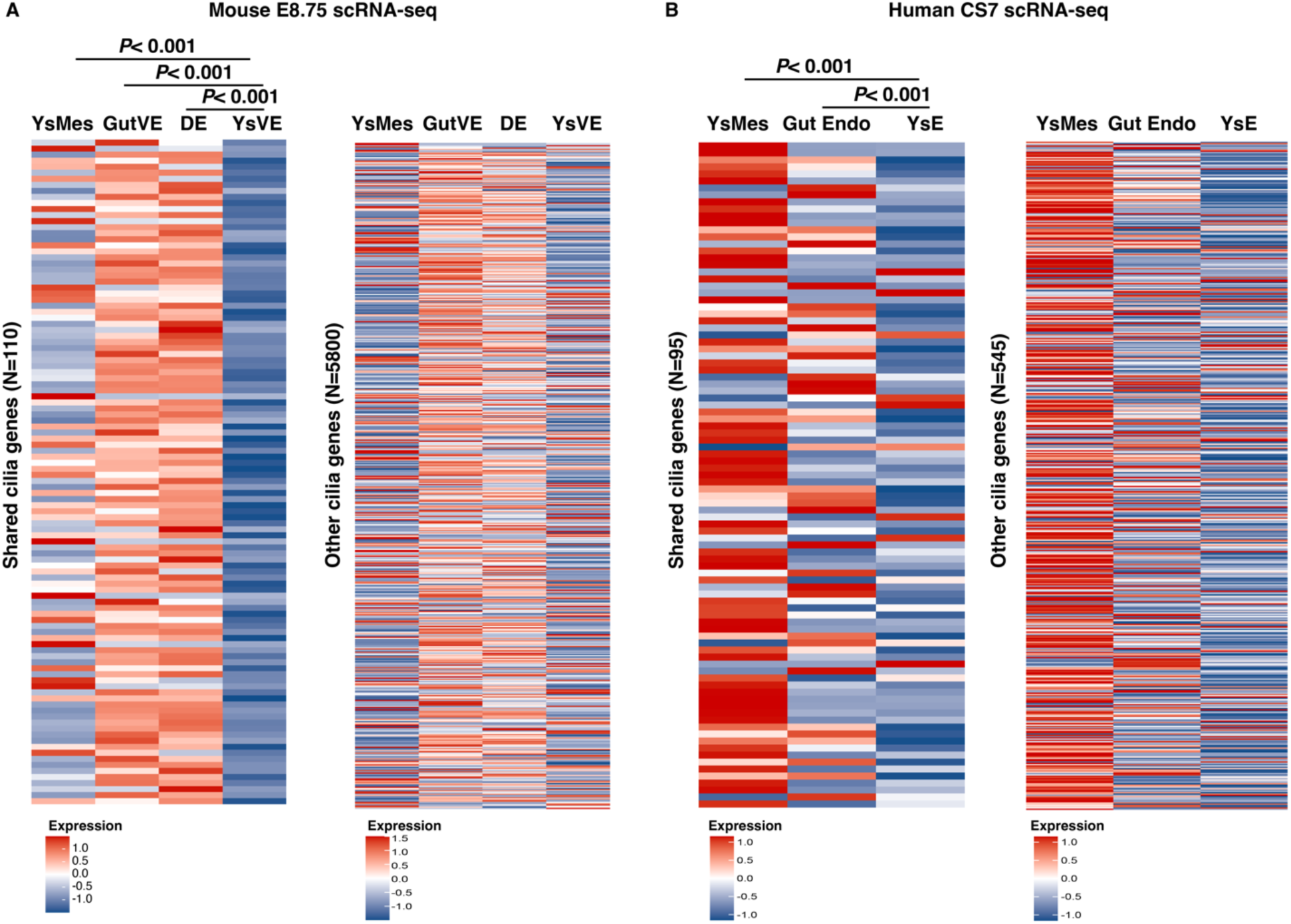
The shared cilia genes are upregulated in ciliated cell types in mouse and human. **(A)** Heatmaps of scRNA-seq expression in mouse E8.75 embryos of the shared cilia genes (N=110, table S10) and the remaining cilia genes (N=580) in YsMes, GutVE, DE compared to YsVE. (**B**) Heatmaps of scRNA-seq expression in human embryos staged CS7 stage (16-19 days post fertilization) of human homologs of the mouse shared cilia genes (N=95) and the remaining cilia genes (N=545) showing general upregulation of only shared genes in yolk sac mesoderm (YsMes) and gut endoderm (Gut Endo) compared to yolk sac endoderm (YsE) (*30*).

**Fig. S4.**
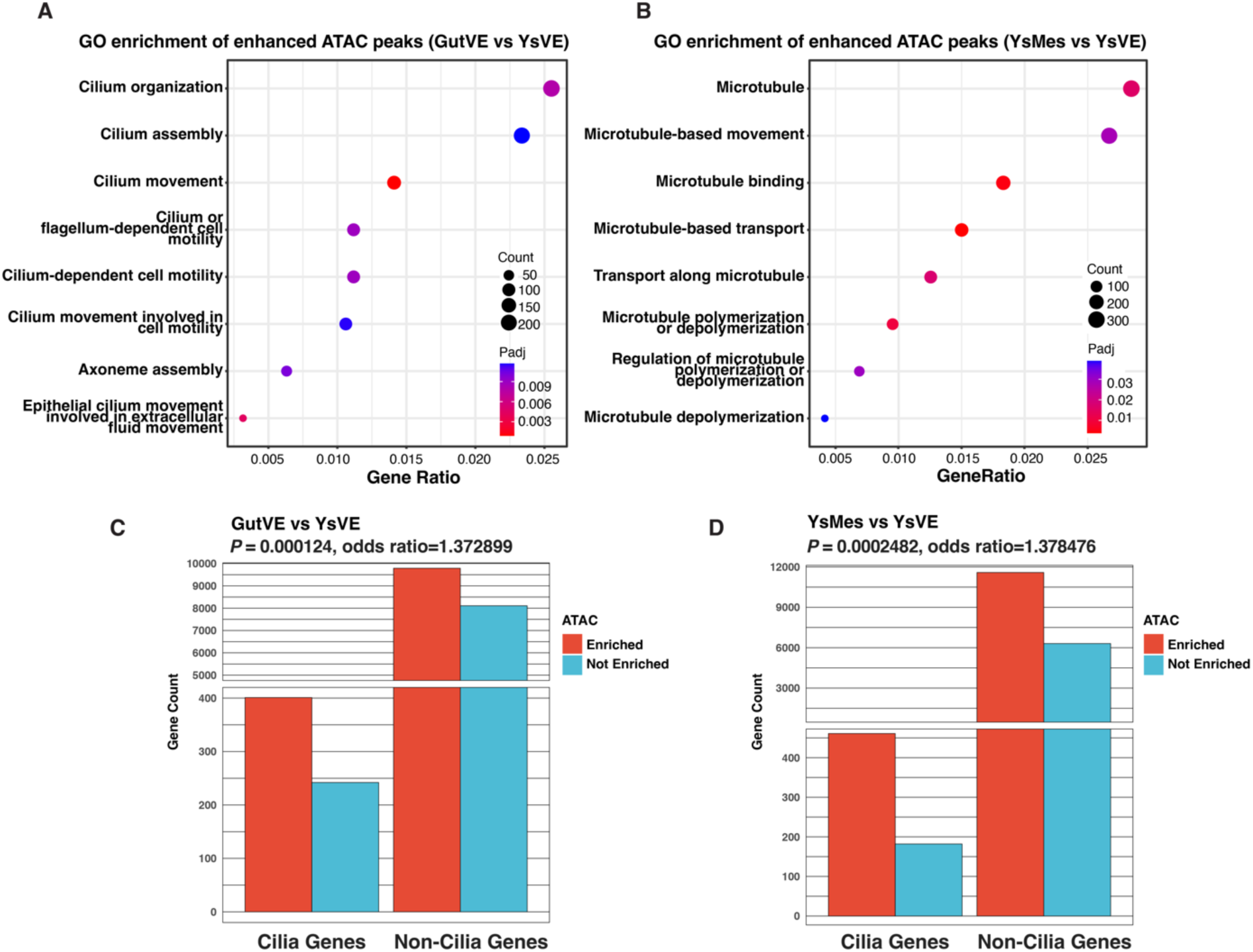
Chromatin regions of cilia genes are more accessible in ciliated cell types than YsVE. **(A**, **B)** Dot plots showing cilia-related biological process terms and microtubule-based transport terms are enriched in genes with enhanced chromatin regions in GutVE (A), and YsMes (B) compared to YsVE at E8.75. (**C**, **D**) Bar plots showing enrichment of cilia genes in enhanced ATAC-seq peaks in GutVE (C) and YsMes (D) compared to YsVE, based on two-sided Fisher’s exact test.

**Fig. S5.**
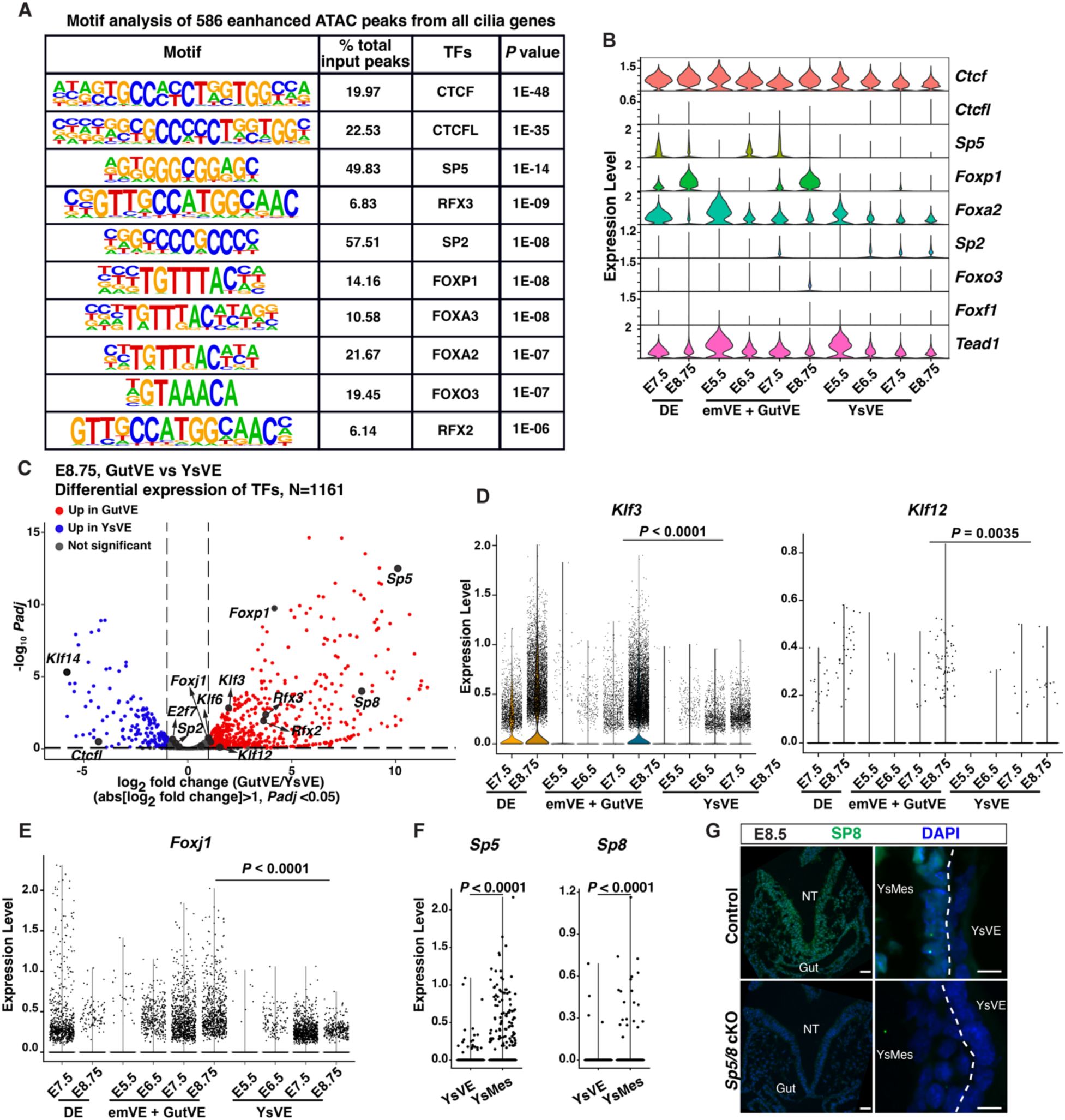
SP5/8 are upregulated in GutVE compared to YsVE. (**A**) Enrichment of transcription factor recognition sequences in 586 differential ATAC-seq peaks of all cilia genes in GutVE and YsMes versus YsVE. The list is ranked based on *P* value. See also table S16. (**B**) Violin plots showing the log_10_ CPM expression levels of the top 10 candidate TFs genes (listed in Fig. 2C) in DE, GutVE, and YsVE at E5.5-E8.75. (**C**) Volcano plot of scRNA-seq data showing differentially expressed genes encoding TFs in GutVE and YsVE at E8.75. The genes listed include the top 10 TFs and additional SP/KLF family and known cilia TFs that are differentially expressed, see also table S17. (**D**, **E**) Violin plots showing the log_10_ CPM expression levels of *Klf3*, *Klf12* and *Foxj1* in DE, emVE+GutVE and YsVE. (**F**) Violin plots showing the log_10_ CPM expression levels of *Sp5*, and *Sp8* in YsVE and YsMes at E8.75. (**G**) IF staining for SP8 on sections of an embryo (left) and yolk sac (right) from Control and *Sp5/8* cKO E8.75 embryos. Scale bars 20μm.

**Fig. S6.**
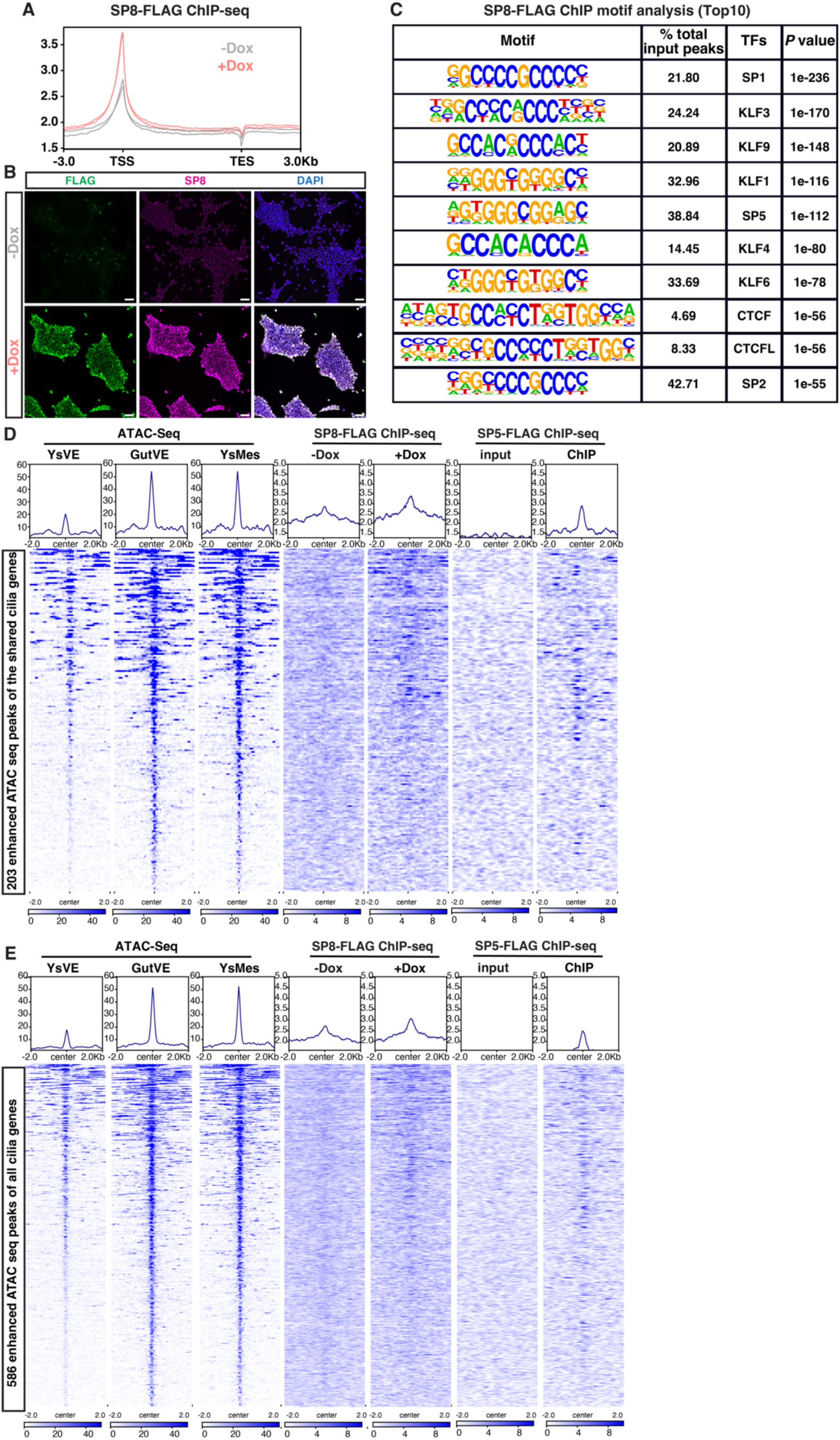
SP5/8 bind cilia genes associated with enhanced chromatin accessibility. (**A**) Composite plots showing distribution of DNA sequences bound by SP8-FLAG in doxycycline-induced (+Dox) and untreated (-Dox) *i-SP8-3xFlag* EpiSCs (N=2 technical replicates/condition). See also table S18. (**B**) IF analysis of FLAG and SP8 in *i*-*Sp8*-3xFLAG EpiSCs with and without Doxycycline treatment. Scale bar 20μm. (**C**) Enrichment of transcription factor recognition sequences in differential SP8-FLAG ChIP-seq peaks (N=10417) identified in +Dox compared to -Dox cells. List is ranked based on *P* value compared to background peaks (N=38135). (**D**, **E**) Tornado plots showing enhanced chromatin regions of the shared cilia genes (D, N=203) and all cilia genes (E, N=586) in ATAC-seq, SP8-FLAG, SP5-FLAG ChIP-seq data.

**Fig. S7.**
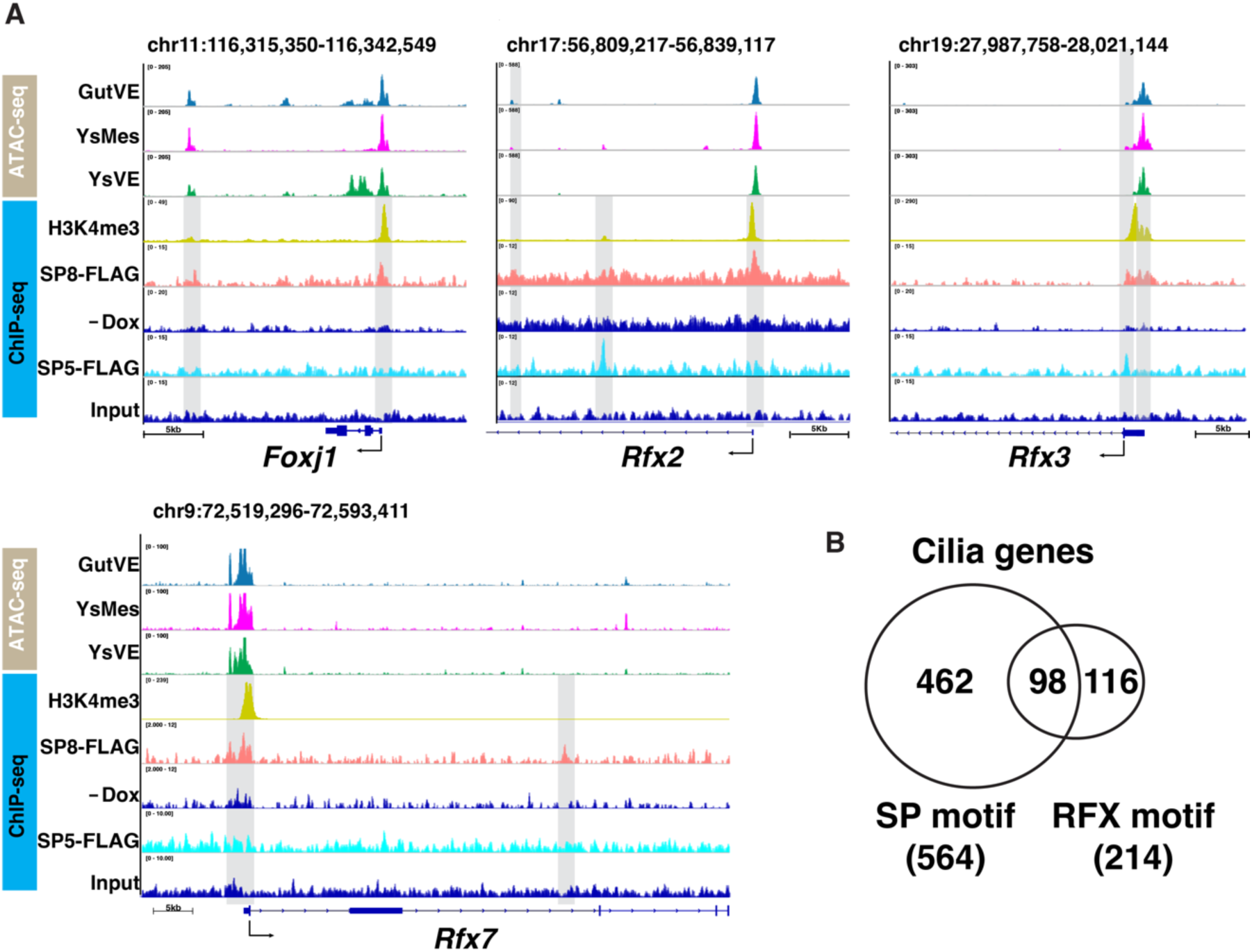
SP5/8 bind *Foxj1* and RFXs TF genes. (**A**) ATAC-seq plots of *Foxj1*, *Rfx2*, *Rfx3,* and *Rfx7* in GutVE, YsMes, YsVE overlapped with ChIP-seq tracks of H3K4me3, SP8-FLAG in EpiSCs and SP5-FLAG in mouse embryoid bodies. SP5/8 binding sites and ATAC-seq peaks with significance enhancement in a single 500 bp peak are highlighted by grey boxes. Y-axis scale was chosen to optimize the visualization of peaks for each sample. (**B**) Venn diagram showing the distribution of SP5 and RFX motifs across all cilia genes.

**Fig. S8.**
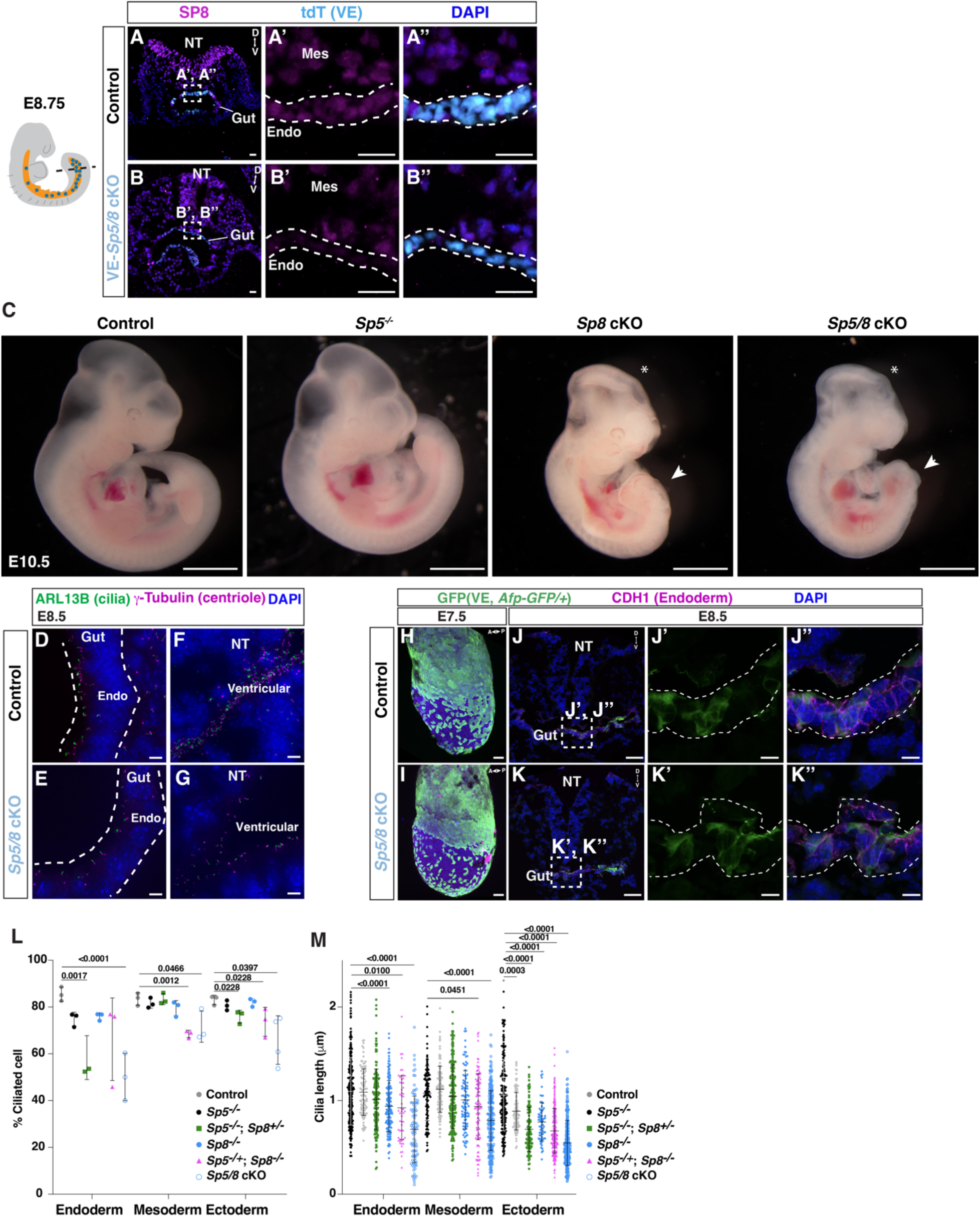
SP5/8 are required for global primary cilia formation. (**A**, **B**) IF staining of transverse sections of gut tube in Control (*Ttr-Cre/+*; *R26^lsl-tdT/+^*; *Sp5^-/+^*; *Sp8^fl/+^*) and *VE*-*Sp5/8* cKO (*Ttr-Cre/+*; *R26^lsl-tdT/+^*; *Sp5^-/-^*; *Sp8^fl/fl^*) E8.75 embryos. NT, neural tube; Mes, mesenchymal cells. The schematic of the embryo was modified from (*21*). (**C**-**F**) IF staining for ARL13B, g-tubulin, and DAPI in transverse sections of gut tube and neural tube in Control and *Sp5/8* cKO embryos (12 somites). Endo, gut endoderm; Mes, mesenchymal cells. (**G**) Bright-field images of Control (*CAG-Cre/+*; *Sp5^-/+^*; *Sp8^fl/+^*), *Sp5^-/-^*, *SP8* cKO (*CAG-Cre/+*; *Sp8^fl/fl^*) and *Sp5/8* cKO (*CAG-Cre/+*; *Sp5^-/-^*; *Sp8^fl/fl^*) embryos at E10.5. Asterisks, brain exencephaly; arrowheads, posterior truncations. (**H**, **I**) Maximum intensity projection of wholemount IF staining in Control (H) and *Sp5/8* cKO (I) embryos, with *Afp-GFP* reporter at E7.5. (**J**, **K**) IF staining of transverse sections of Control (J) and *Sp5/8* cKO (K) embryos with *Afp-GFP/+*allele. (**L**, **M**) Quantification of the percentage of ciliated cells (L) and cilia length (M) in Control, *Sp5^-/-^*, *Sp5^-/-^*; *Sp8^-/+^*, *Sp8^-/-^*, *Sp5^-/+^*; *Sp8^-/-^* and *Sp5/8* cKO gut endoderm, mesoderm (mesenchymal) and neural ectoderm cells at E8.5 (N≥3 embryos/genotype). A, anterior; P, posterior; D, dorsal; V, ventral. Scale bar, 50μm (A, B, H, I), low magnification, 10μm insets (C, D, J, K). Statistical analysis: one-way ANOVA. (L, M).

**Fig. S9.**
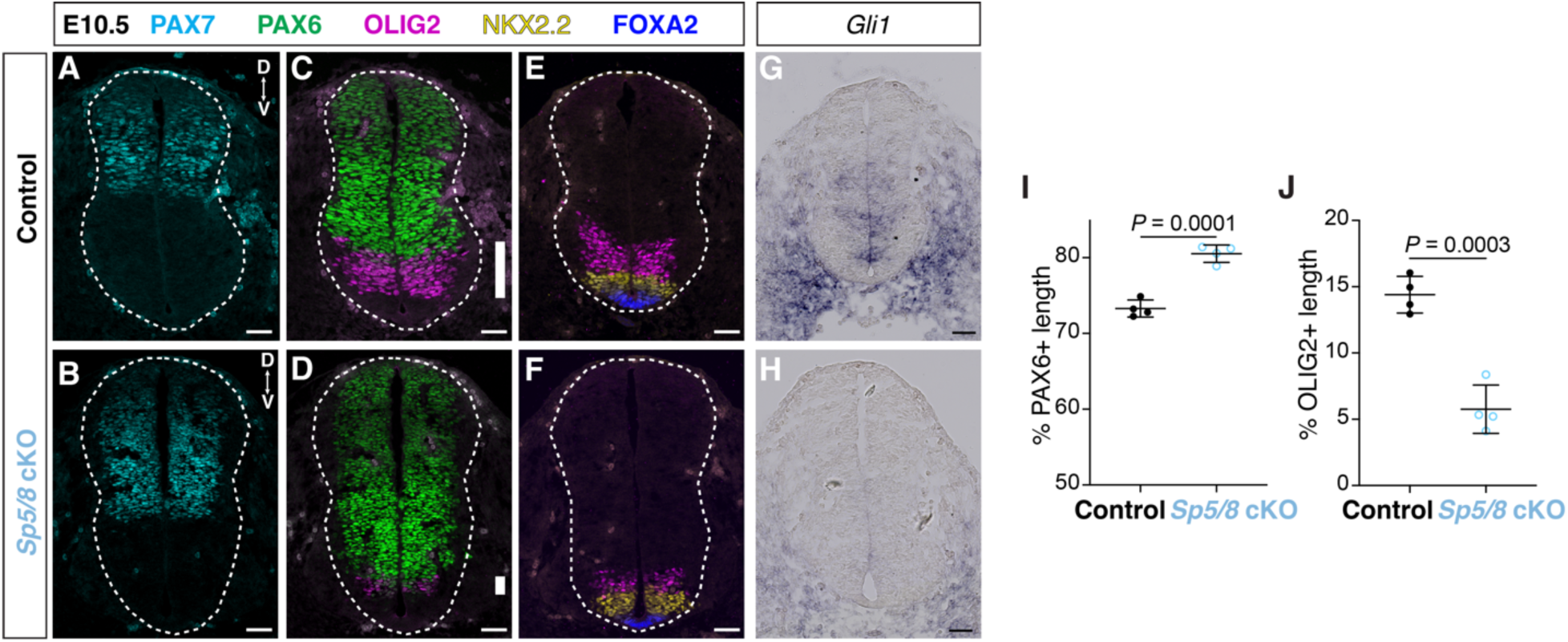
*Sp5/8* cKO mutant have a reduction of HH signaling. (**A**-**F**) IF staining of lumbar neural tube sections in Control and *Sp5/8* cKO embryos (E10.5), with SHH-dependent ventral progenitor types marked by FOXA2 (floor plate), NKX2.2 (V3 interneuron), OLIG2 (motor neuron), and PAX6 (dorsal) (N=4 embryos per genotype). White bar indicates the range of OLIG2^+^ nucleus (C, D). (**G**, **H**) RNA in situ hybridization of *Gli1* in transverse sections of lumbar neural tube in Control and *Sp5/8* cKO embryos (E10.5, N=4 per genotype). (**I**, **J**) Quantification of the percentage length of the PAX6^+^ (I) and OLIG2^+^ (J) domains of the neural tube. Scale bar, 50μm (A-H). Statistical analysis: unpaired *t*-test (I, J).

**Fig. S10.**
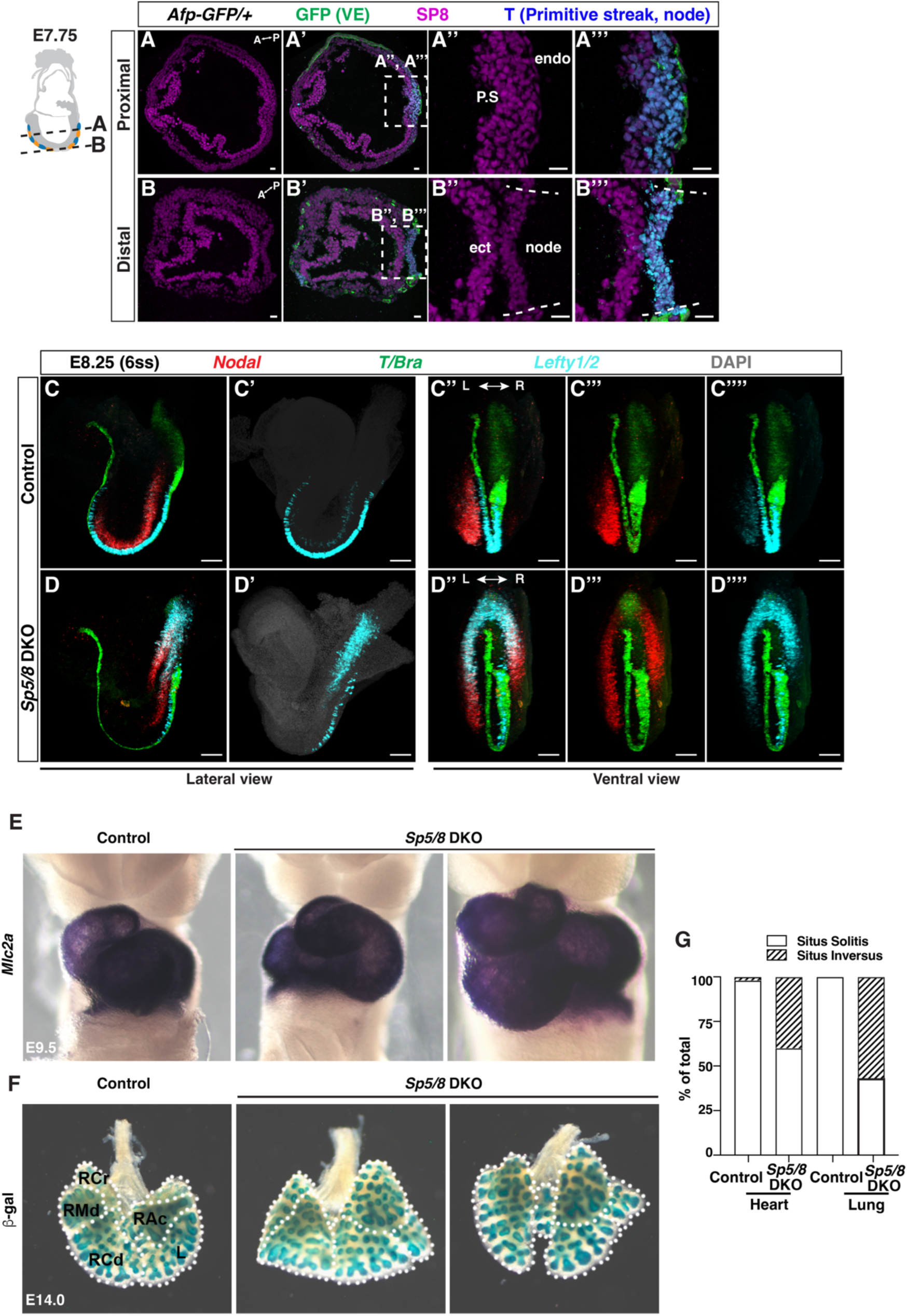
SP5/8 regulate left-right asymmetry of organs. (**A**, **B**) IF analysis of sections in an E7.75 *Afp-GFP/+* embryo for SP8 and the node/primitive streak marker (T) in the proximal (A) and distal embryo (B) embryo (node is distal). P.S, primitive streak; ect, ectoderm; endo, endoderm. A, anterior; P, posterior. Scale bar, 20μm. The schematic of the embryo was modified from (*21*). (**C**, **D**) Maximum intensity projection images of hybridization chain reaction in situ analysis of littermate Controls (*Sp5^-/-^*; *Sp8^-/+^*, 6-7 somites) and *Sp5/8* DKO (*Sp5^-/-^*; *Sp8^-/-^*, 5-6 somites). Scale bar, 100μm. (**E**) Wholemount RNA in situ hybridization for Myosin regulator light chain 2 (*Mlc2a*) in E9.5 littermate Control (*Sp5^-/-^*; *Sp8^-/+^*) and *Sp5/8* DKO (*Sp5^-/-^*; *Sp8^-/-^*) embryos (N=3/genotype). (**F**) X-Gal staining of E14.0 Control and *Sp5/8* DKO embryos (N=3/genotype). X-Gal staining detects ß-Gal protein in the *Sp5 lacZ* knock-in null allele. RCr, Right Cranial; RMd, Right Middle; RCd, Right Caudal; RAc, Right Accessory; L, Left lobe. **G**) Bar plot showing organ situs phenotype in Control (*Sp5^-/-^*; *Sp8^+/+^* or *Sp5^-/-^*; *Sp8^-/+^*) and *Sp5/8* DKO embryos at E9.5 (heart, N=47, 20) and E14.0 (lung, N=12, 7).

**Fig. S11.**
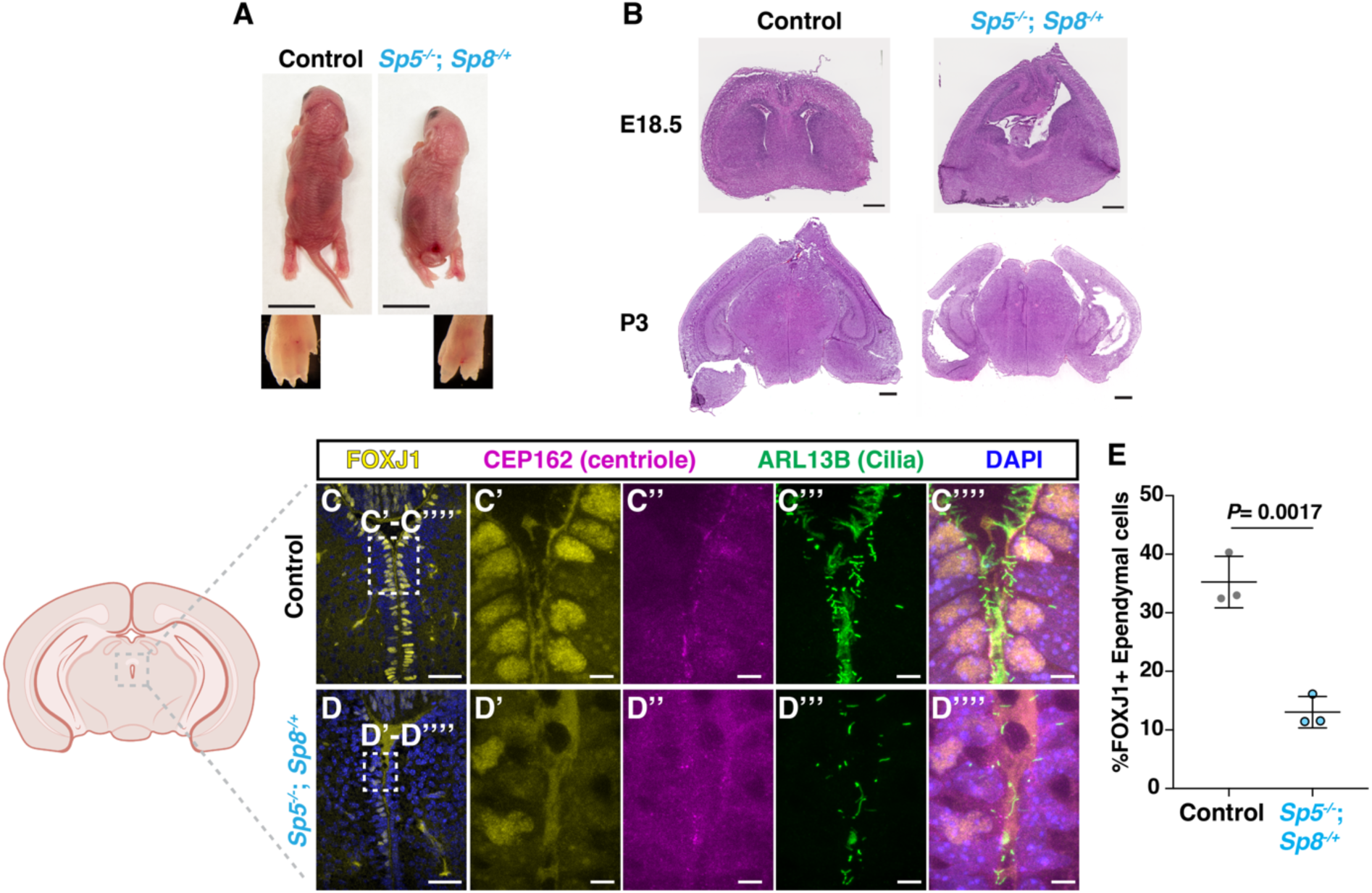
SP5/8 regulate motile cilia formation. (**A**) Bright field images of Control (*Sp5^-/+^*) and *Sp5^-/-^*; *Sp8^-/+^* pups at P0. High magnification shows limb patterning defect. (B) Hematoxylin and Eosin staining of control and *Sp5^-/-^*; *Sp8^-/+^*brain coronal sections at P0 and P3 (N≥3 per genotype). (**C, D**) IF staining of motile cilia markers and FOXJ1 in the third ventricular (as indicated in the schematic) in Control and *Sp5^-/-^*; *Sp8^-/+^*brains at P0. Schematic created with BioRender.com. (**E**) Quantification of the percentage of FOXJ1^+^ cells of total ependymal cells in the third ventricular (N=3 per genotype). Scale bar 0.5mm (A, B); 20μm (C, D), insets 5μm. Statistical analysis: unpaired *t*-test (E).

**Fig. S12.**
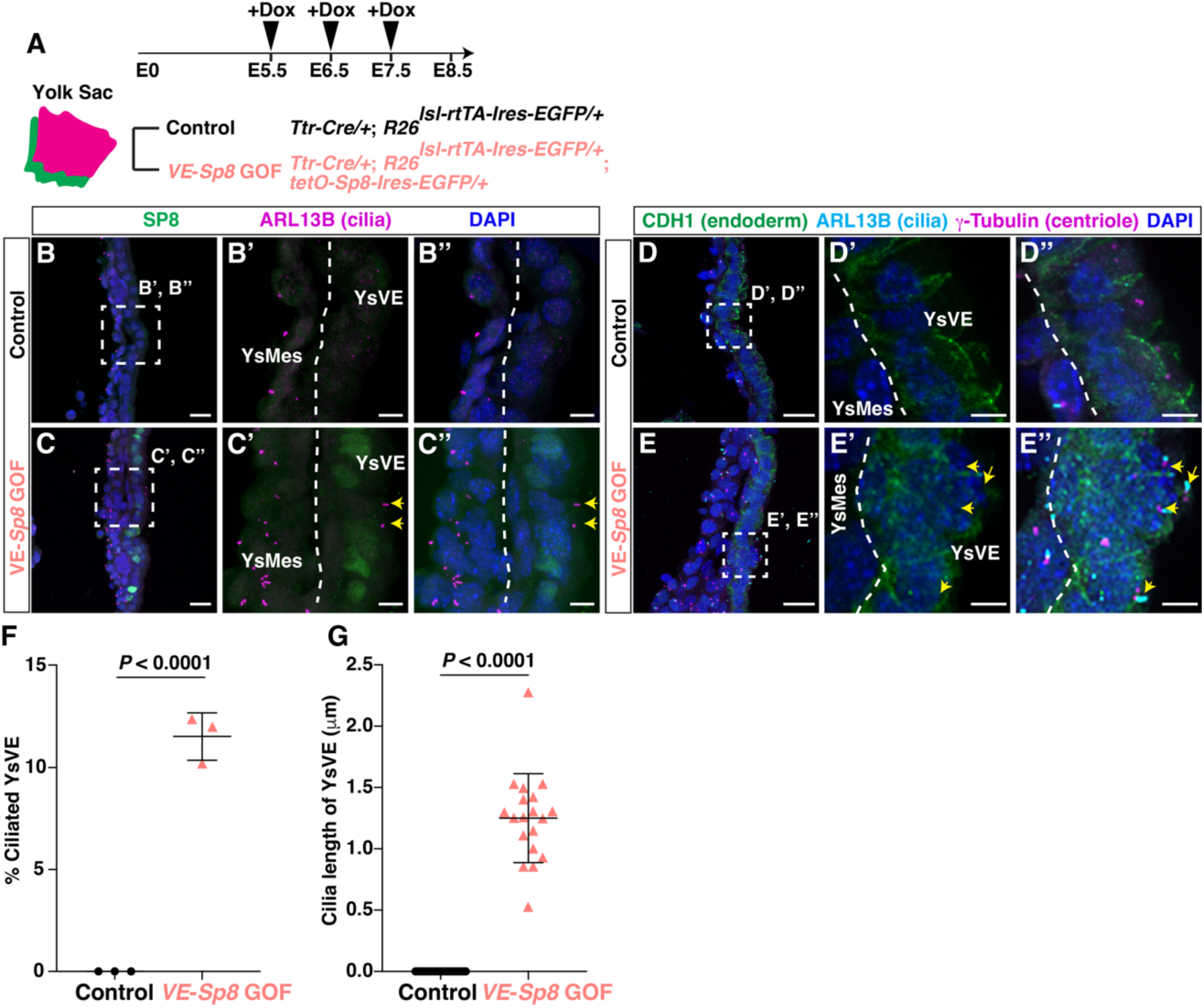
SP8 mis-expression induces cilia in YsVE. (**A**) Schematic of *VE-Sp8* gain-of-function (GOF) strategy. (**B**, **C**) IF staining of yolk sac sections from Control (B) and *VE*-*Sp8* GOF (C) embryos for SP8, ARL13B and DAPI. Dashed line, separation between YsVE and YsMes. (**D**, **E**) IF staining of yolk sac sections from Control (D) and *VE*-*Sp8* GOF (E) embryos for CDH1, ARL13B, g-Tubulin and DAPI. (**F**, **G**) Quantification of the percentage of ciliated cells (F) and cilia length (G) in YsVE in Control and *VE*-*Sp8* GOF embryos (N=3 embryos/genotype). Statistical analysis: unpaired *t*-test. Scale bars, 20μm (B-E).

**Fig. S13.**
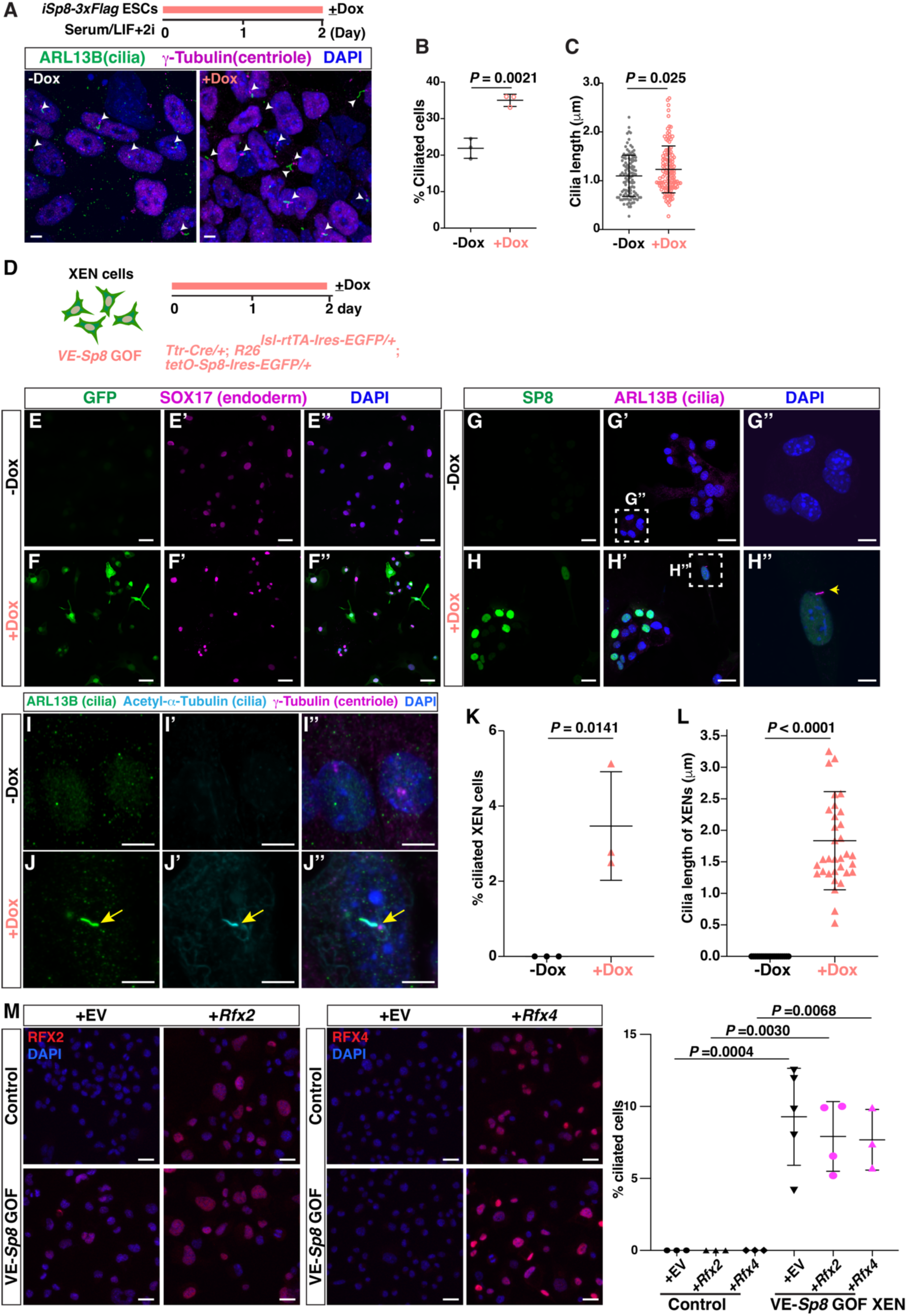
SP8 mis-expression induces cilia in XEN. (**A**) IF staining in *i-Sp8-3xFlag* ESCs ±Dox for 48 hours. (**B**, **C**) Quantification of the percentage of ciliated cells (B) and cilia length (C) in *i-Sp8-3xFlag* ESCs ±Dox (N=3 technical replicates per condition). (**D**) Schematic of inducible *VE*-*Sp8* GOF strategy. (**E**-**J**) IF staining of inducible *VE*- *SP8* GOF XEN cells ±Dox for 48 hours. (**K**, **L**) Quantification of the percentage of ciliated cells (K) and cilia length (L) in Control and *VE*-*Sp8* GOF XEN cells (N=3 cell lines, ±Dox). Dashed line squares indicate high power image to right; yellow arrows, ciliated XEN cells. (**M**) Quantification of the percentage of ciliated cells in Control cells (-Dox) and *VE*-*Sp8* GOF XEN cells (+Dox for 48 hours) that had been infected with dox-inducible lentivirus expressing RFX2 or RFX4 proteins with turbo RFP or only turbo RFP (EV), N=3 technical replicates per condition. Scale bars, 20μm (B-E, M) and 5μm (G’’, H’’, I, J). Statistical analysis: unpaired *t*-test (K, L) or two-way ANOVA (M).

**Fig. S14.**
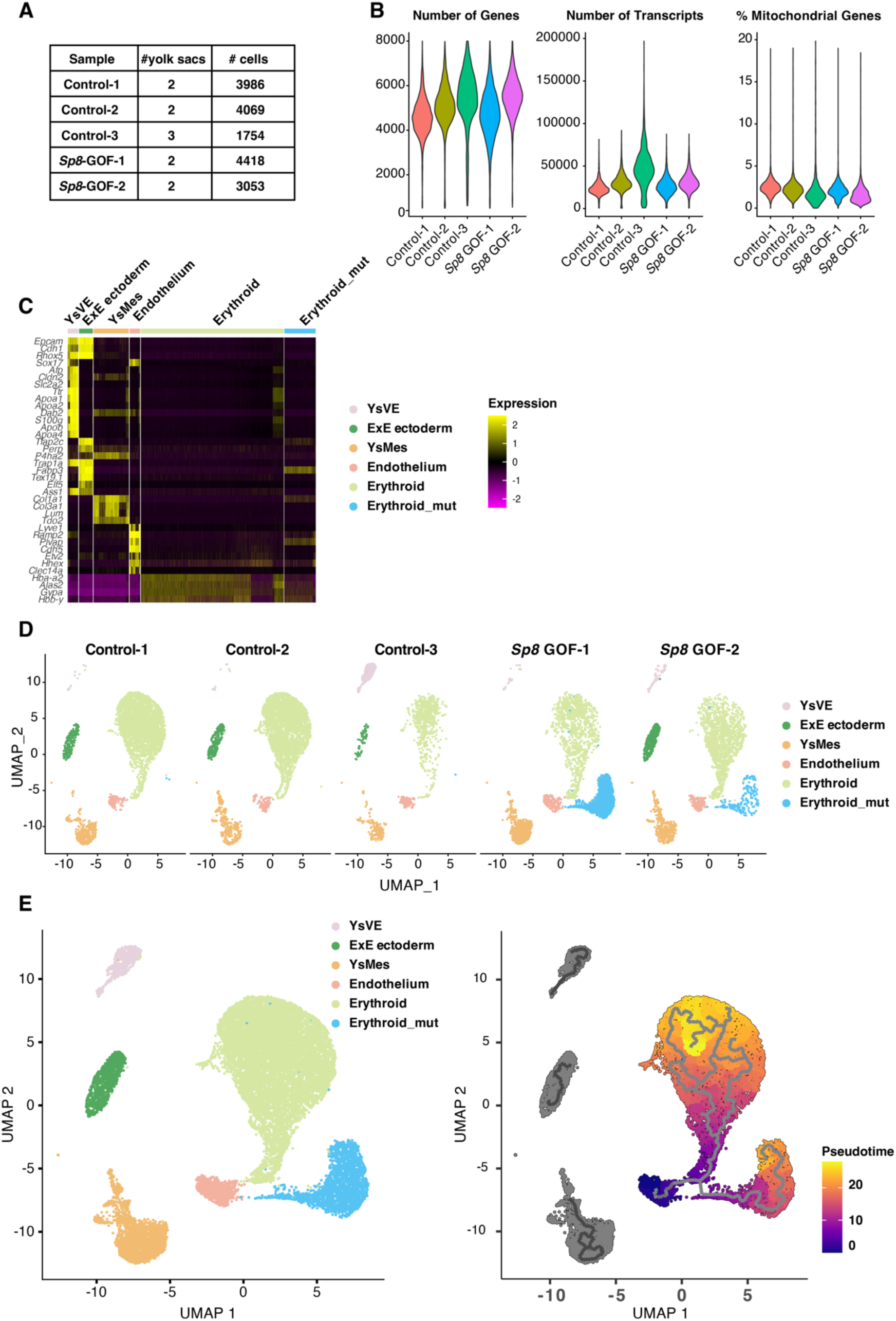
Quality control and clustering analysis of *Sp8* GOF yolk sac scRNA-seq analysis. **(A)** Table summarizing the number of yolk sacs and corresponding cell counts per sample. **(B)** Violin plots depicting number of genes, number of transcripts and percentage of mitochondrial genes. (**C**) Heatmap plot displaying marker gene expression across different clusters. (**D**) UMAP plots showing the cluster distribution of cells from each sample. Control samples: Control-1 and Control-2 each have two yolk sac from E8.5 embryos, Control-3 is the cells from combined 3 yolk sacs into one sample. *Sp8* GOF samples: each contains two E8.5 yolk sacs. (**E**) UMAP plots showing the clusters of cells with all samples combined (left) and a pseudotime trajectory of the erythroid lineage (right).

**Fig. S15.**
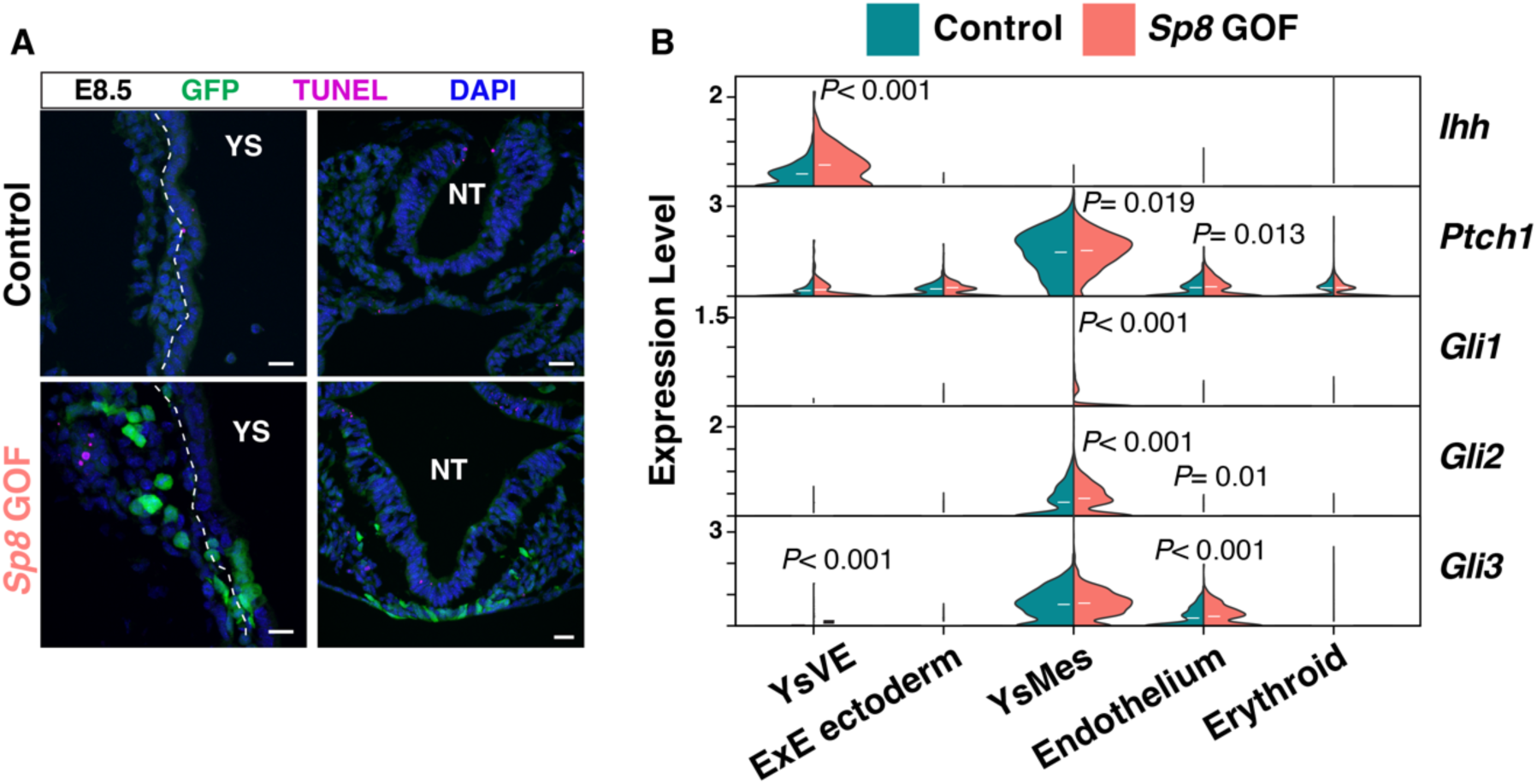
Cell death and HH signaling activity are not altered in *Sp8* GOF embryos. (**A**) TUNEL (terminal deoxynucleotidyl transferase dUTP nick end labeling) staining of sections of yolk sac and neural tube from Control and *Sp8* GOF embryos at E8.5. YS, yolk sac. NT, neural tube. Scale bars, 20μm. (**B**) Violin plots showing the expression of HH signaling pathway components across different cell types in Control and *Sp8* GOF yolk sac cell types (clusters). Statistical analysis: Wilcoxon rank-sum tests.

**Fig. S16.**
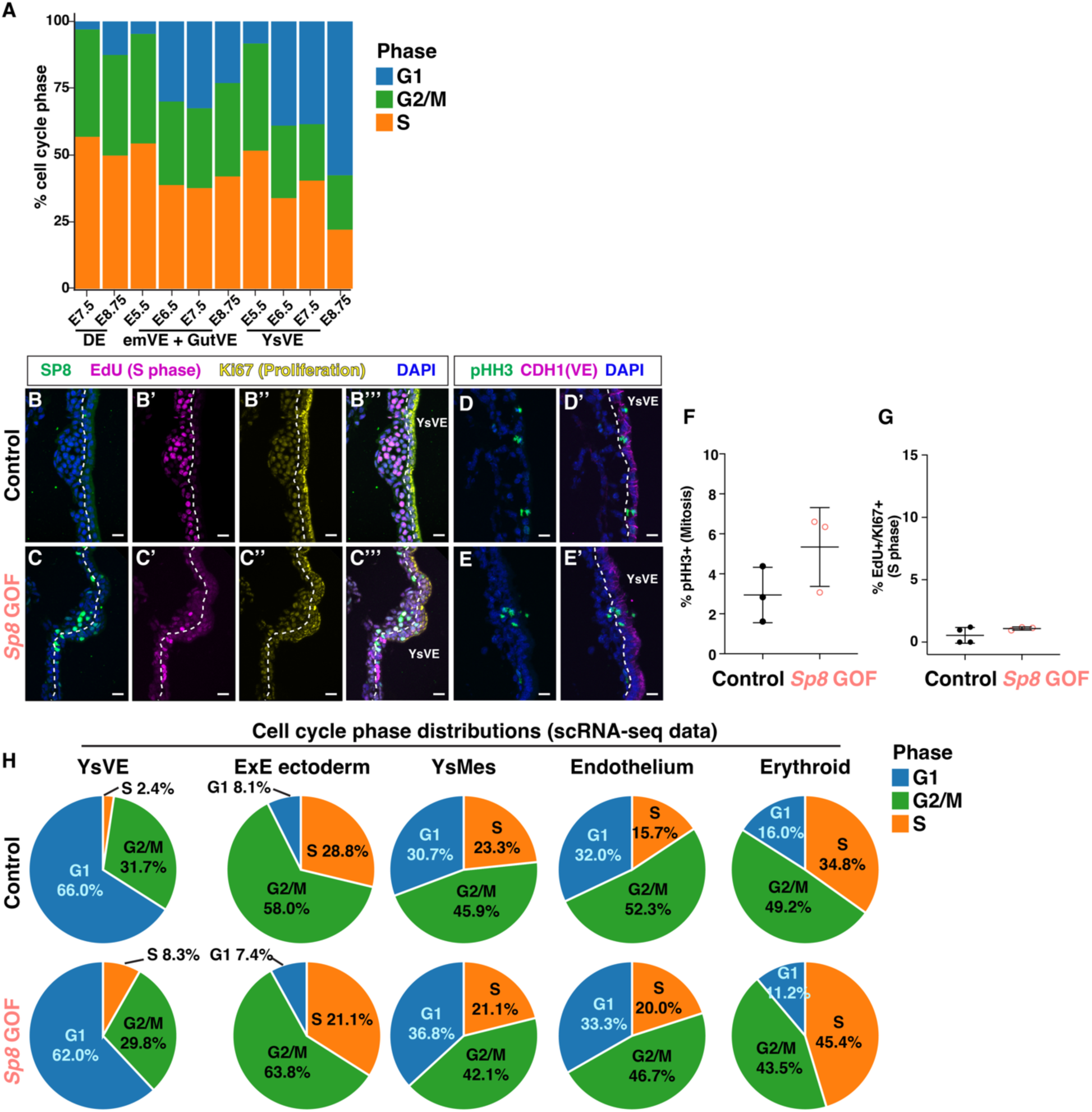
SP8 over-expression does not extend G1/G0 phase in the yolk sac. (**A**) Stacked bar plots showing the distribution of cell cycle phase in DE, GutVE, and YsVE at E5.5-E8.75 using scRNA-seq data. **(B**-**E)** IF staining of cell cycle markers, pHH3 (mitosis marker), EdU (S phase marker) and KI67 (proliferation marker), in transverse section of yolk sacs from Control and *Sp5/8* cKO embryos at E8.5. Dashed lines indicate the separation between YsVE and other cells. Scale bars, 20μm. **(F**, **G)** Quantification of the percentage of pHH3^+^ (F) and EdU⁺ KI67⁺ of KI67^+^ (G) cells in YsVE of Control and *Sp5/8* cKO embryos (N≥3 embryos/condition). Statistical analysis: unpaired *t*-test. (**H**) Cell cycle phase distribution across cell types in Control and *Sp8* GOF yolk sac tissue based on scRNA-seq data.

**Fig. S17.**
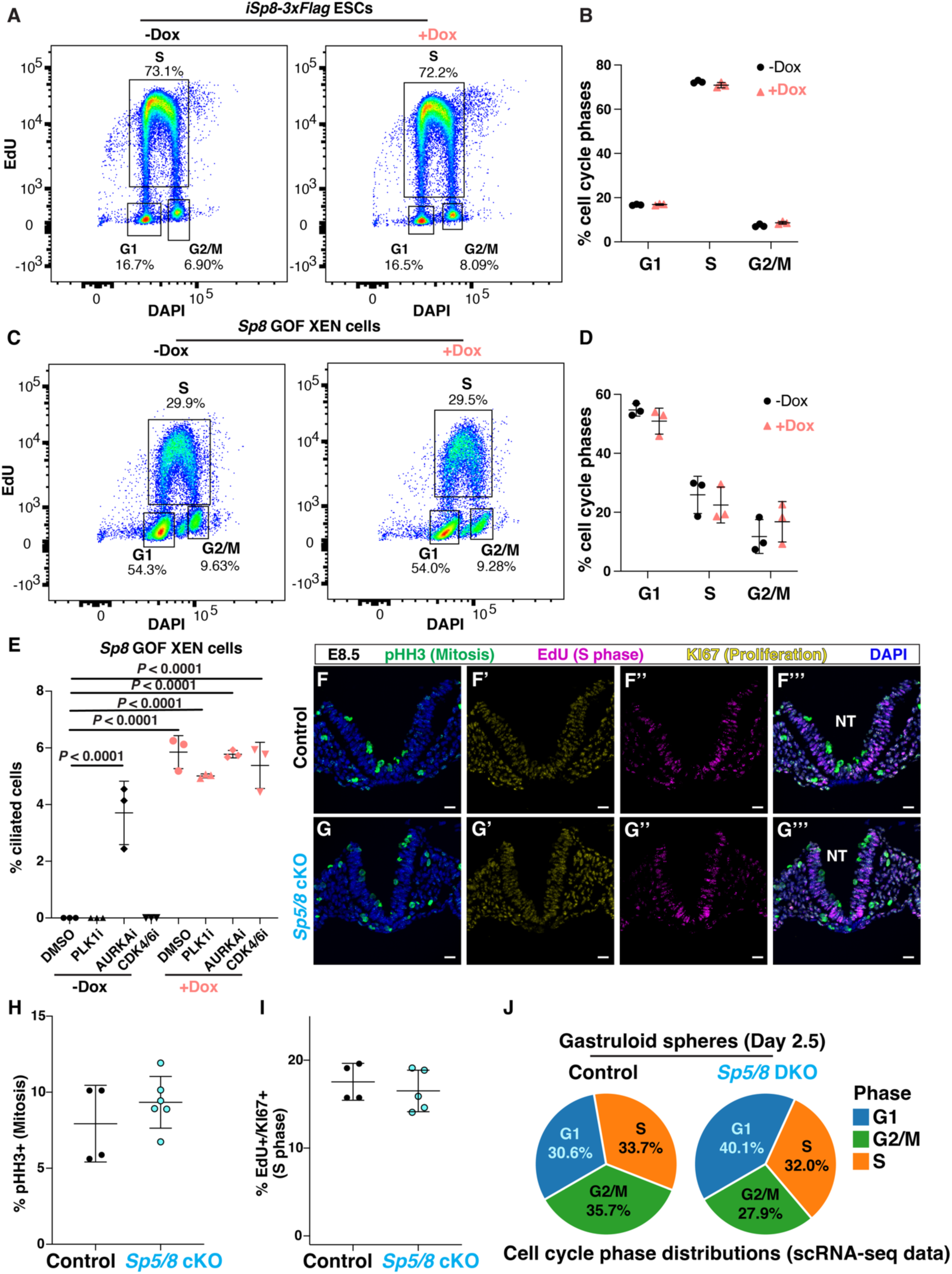
SP8 promotes cilia formation independently of cell cycle regulation. (**A**) Flow cytometry analysis of DAPI and EdU-Cy5 staining in *i-Sp8-3xFlag* ESCs ±Dox for 48 hours. (**B**) Quantification of cell cycle phase distribution from (A), N=3 technical replicates/condition. (**C**) Flow cytometry analysis of DAPI and EdU-Cy5 staining in *VE*-*Sp8* GOF XEN ±Dox 48 hours. (**D**) Quantification of cell cycle phase distribution from (C), N=3 technical replicates per condition. (**E**) Quantification of the percentage of ciliated cells in *VE*-*SP8* GOF XEN ±Dox for 48 hours and treated with DMSO (vehicle), PLK1i (BI2536, 1µM), AURKAi (250nM), CDK4/6i (Palbociclib, 250nM) to arrest cell cycle at G2/M, G1/G0 for 16 hours and AURKAi also inhibits proteins that disassemble cilia. N=3 technical replicates/condition. (**F**, **G**) IF staining of cell cycle markers, pHH3 (mitosis), EdU (S phase) and KI67 (proliferation), in transverse sections of neural tube region from Control (F) and *Sp5/8* cKO embryos (G) at E8.5. Scale bars, 20μm. **(H**, **I)** Quantification of the percentage of pHH3^+^ (H) and EdU⁺ KI67⁺ of KI67^+^ (I) cells in the neural ectoderm of Control and *Sp5/8* cKO embryos. Statistical analysis: two-way ANOVA (E) or unpaired *t*-test (H, I). (**J**) Cell cycle phase distribution across cell types in Control and *Sp5/8* DKO gastruloids based on scRNA-seq data.

**Table S1 to S25.**

**Table S1. List of Syscilia cilia genes (N=690)** (***7***).

**Table S2. Gene ontology terms (biological processes) based on genes upregulated in GutVE compared to YsVE at E7.5 (*Padj*<0.05). (related to Fig. 1I)**

**Table S3. Gene ontology terms (biological processes) based on genes upregulated in GutVE compared to YsVE at E8.75 (*Padj*<0.05). (related to Fig. 1J)**

**Table S4. scRNA-seq differential gene expression analysis of cilia genes (N=690) in GutVE compared with YsVE at E5.5-8.75. (*Padj*<0.05). (related to fig. S2A, B)**

**Table S5. Top cell markers for each cluster in yolk sac scRNA-seq data. (related to fig. S2C)**

**Table S6. Gene ontology terms (biological processes) based on genes upregulated in YsMes compared to YsVE at E8.75 (*Padj*<0.05). (related to fig. S2D)**

**Table S7. scRNA-seq differential gene expression analysis of cilia genes (N=690) in YsMes compared with YsVE at E8.75 (*Padj*<0.05). (related to fig. S2E)**

**Table S8. Gene ontology terms (biological processes) based on genes upregulated in DE compared to YsVE at E8.75 (*Padj*<0.05). (related to fig. S2F)**

**Table S9. scRNA-seq differential gene expression analysis of cilia genes (N=690) in DE compared with YsVE at E8.75 (*Padj*<0.05). (related to fig. S2G)**

**Table S10. scRNA-seq analysis showing shared cilia genes upregulated in GutVE, DE and YsMes compared to YsVE at E8.75 (log2FC>1, *Padj*<0.05). (related to Fig. 1K)**

**Table S11. Gene ontology terms (biological processes) based on ATAC peaks/genes upregulated in GutVE compared to YsVE at E8.75 (*Padj*<0.05). (related to fig. S4A)**

**Table S12. Gene ontology terms (biological processes) based on peaks/genes upregulated in YsMes compared to YsVE at E8.75 (*Padj*<0.05). (related to fig. S4B)**

**Table S13. ATAC-seq analysis listing regions in Syscilia cilia genes with enhanced open chromatin in GutVE and YsMes compared to YsVE (log2FC>0, *Padj*<0.05). (related to Fig. 2A)**

**Table S14. 203 regions with enhanced open chromatin based on ATAC-seq analysis of the core ciliome genes in GutVE and YsMes compared to YsVE (log2FC>0, *Padj*<0.05). (related to Fig. 2C)**

**Table S15. Enrichment of transcription factor recognition sequences in 203 enhanced ATAC peaks in the shared cilia genes in GutVE and YsMes compared to YsVE (q-value<0.05). (related to Fig. 2C)**

**Table S16. Enrichment of transcription factor recognition sequences in 586 enhanced ATAC peaks in all cilia genes in GutVE and YsMes compared to YsVE (q-value<0.05). (related to fig. S5A)**

**Table S17. scRNA-seq analysis showing expression of known TFs (N=1161) in GutVE compared to YsVE at E8.75 (*Padj*<0.05). (related to fig. S5C)**

**Table S18. Peak calling results for cilia genes from SP5-FLAG ChIP-seq data analysis based on Dox-induced versus input i-*Sp5*-3xFlag embryoid bodies (*P*<0.1). (related to Fig. 3B)**

**Table S19. Peak calling results from SP8-FLAG ChIP-seq data analysis based on Dox-induced versus untreated i-*Sp8*-3xFlag EpiSC (*P*<0.01). (related to fig. S6A, C)**

**Table S20. Peak calling results for cilia genes from SP8-FLAG ChIP-seq data analysis based on Dox-induced versus untreated i-Sp8-3xFlag EpiSC (*P*<0.01). (related to Fig. 3B)**

**Table S21. Differential gene expression of SP5/8 bound cilia genes in the three E8.75 ciliated cell types vs YsVE. (related to Fig. 3C)**

**Table S22. scRNA-seq differential gene expression analysis of SP5/8 bound cilia genes (N=187) in *Sp5/8* DKO compared to control gastruloids (*Padj*<0.05). (related to Fig. 3K)**

**Table S23. Gene ontology analysis of differentially expressed genes downregulated in *Sp5/8* DKO compared to control gastruloids (*Padj*<0.05). (related to Fig. 3L)**

**Table S24. scRNA-seq differential gene expression analysis of SP5/8 bound cilia genes (N=187) in *Sp8* GOF compared to control YsVE. (related to Fig. 5G)**

**Table S25. Reagents and primer information.**

## Notes

### Competing Interest Statement

The authors have declared no competing interest.

